# Detection and quantification of the vacuolar H^+^-ATPase using the *Legionella* effector protein SidK

**DOI:** 10.1101/2021.07.29.454369

**Authors:** Michelle E. Maxson, Yazan M. Abbas, Jing Ze Wu, Sergio Grinstein, John L. Rubinstein

## Abstract

Acidification of secretory and endocytic organelles is required for proper receptor recycling, membrane traffic, protein degradation, and solute transport. Proton-pumping vacuolar ATPases (V-ATPases) are responsible for this luminal acidification, which increases progressively as secretory and endocytic vesicles mature. An increasing density of V-ATPase complexes is thought to account for the gradual decrease in pH, but available reagents have not been sufficiently sensitive nor specific to test this hypothesis. We introduce a new probe to localize and quantify V-ATPases in eukaryotic cells. The probe is derived from SidK, a *Legionella pneumophila* effector protein that binds to the V-ATPase A subunit. We generated plasmids encoding fluorescent chimeras of SidK_1-278_, and labeled recombinant SidK_1-278_ with AlexaFluor-568 to visualize and quantify V-ATPases with high specificity in live and fixed cells, respectively. We show that V-ATPases are acquired progressively during phagosome maturation, that they distribute in discrete membrane subdomains, and that their density in lysosomes depends on the subcellular localization of the lysosome.

## INTRODUCTION

The steady-state pH of individual cellular compartments is a key determinant of their function and must be regulated stringently. Indeed, the luminal pH directs receptor recycling and membrane traffic, regulates protein degradation, and contributes to the transmembrane protonmotive force that drives the transport of a variety of organic and inorganic solutes (Maxfield and McGraw, 2004; Saftig and Klumperman, 2009; Fisher and Scheller, 1988; Huotari and Helenius, 2011). It is therefore not surprising that dysregulation of organellar pH has been implicated in various human diseases such as cancer, neurological disorders, osteoporosis, and autoimmunity (Marshansky et al., 2014; Capecci and Forgac, 2013; Colacurcio and Nixon, 2016; Sun-Wada et al., 2006; Eaton et al., 2021).

Most endocytic and secretory organelles maintain an acidic lumen, with acidification increasing progressively as these organelles approach their terminal stages (Mellman et al., 1986). Thus, the endocytic pathway progresses from slightly acidic early endosomes (pH 6.5) to highly acidic lysosomes (pH 4 to 5). The main driver of organellar acidification is the ATP-dependent proton pump known as the vacuolar H^+^-ATPase (V-ATPase). The V-ATPase is present in vesicular membranes of the endocytic and secretory pathways, and is also found in the plasma membrane of specialized cell types involved in the active extrusion of cytosolic protons, such as osteoclasts and renal intercalated cells (Toei et al., 2010; Futai et al., 2019). The V-ATPase is a large rotary complex with 16 different subunits in mammals. ATP-hydrolysis occurs in the soluble catalytic V_1_ region (subunits A to H), driving rotation of the enzyme’s rotor subcomplex and proton translocation through the V_O_ regions (subunits a, d, e, f c, c□, ATP6AP1/Ac45, and ATP6AP2/PRR; Abbas et al., 2020). V-ATPase activity is regulated by several mechanisms, including reversible dissociation of the V_1_ and V_O_ regions (Tabke et al., 2014; Kawasaki-Nishi, 2001; Parra and Kane, 1998; Poëa-Guyon et al., 2013), phosphorylation (Voss et al., 2007; Alzamora et al., 2010), and changes in membrane lipid composition (Banerjee et al., 2019; Vasanthakumar et al., 2019; Uchida et al., 1985).

Several parameters determine the steady-state pH of the lumen of an organelle. Because proton pumping by V-ATPase is electrogenic, the rate of pumping can be limited by the permeability of the membrane to neutralizing counter ions. In addition, the accumulation of protons is opposed by ongoing proton backflux or “leak” via a collection of incompletely characterized channels and transporters. A final fundamental parameter is the density of V-ATPase complexes in the membrane of any particular organelle. It has been tacitly assumed that V-ATPase density increases progressively as the components of endocytic and secretory pathways mature and become more acidic. However, this assumption has not been validated for two main reasons. First, it is difficult to isolate individual stages of these pathways with sufficient purity for reliable biochemical analysis. Second, reagents with sufficient resolution and accuracy are lacking for localization and quantification of V-ATPase in cells. Several antibodies to different V-ATPase subunits are available commercially and some of them have yielded satisfactory results in histological analyses, especially of tissues like the kidney where specialized cell types are uniquely enriched in V-ATPases. However, these same antibodies show poor specificity and a low signal-to-noise ratio when used to stain non-specialized single cells, which have a lower abundance of V-ATPases. The interpretation of the resulting immunostaining can be ambiguous, confounding the results.

The paucity and unsatisfactory performance of reagents currently available to study the V-ATPase motivated us to develop a novel tool for its specific intracellular labeling in eukaryotic cells. To this end we took advantage of SidK, an effector protein deployed by *Legionella pneumophila* to inhibit the V-ATPase (Xu et al., 2010). SidK was recently shown to bind directly to yeast and mammalian V-ATPases by (Zhao et al., 2017; Abbas et al., 2020), allowing use of the effector to purify proton pumps from tissue extracts (Abbas et al., 2020). In this study, we describe the generation and labeling of a recombinant fragment of SidK and its use to localize and quantify V-ATPases in eukaryotic cells with high sensitivity and specificity. We utilized this reagent to study the distribution of the V-ATPase in various cell types, and to monitor its acquisition by membrane-bound compartments as they mature along the endocytic pathway. Lastly, we used SidK to estimate the number of V-ATPase complexes in individual compartments as a function of their position within the cell.

## RESULTS

### Generation of *fluorescent* SidK chimeras for expression in mammalian cells

SidK was reported to bind with high affinity to the A subunit of the V-ATPase (Xu et al., 2010; Sharma and Wilkens, 2017; Abbas et al., 2020; Zhao et al., 2017). We reasoned that the *Legionella* effector would be an effective probe to visualize V-ATPase by fluorescence microscopy. To this end we generated chimeric constructs consisting of amino acids 1 to 278 of SidK (SidK_1-278_) attached to a fluorescent protein (GFP or mCherry) with a linker sequence. SidK_1-278_ suffices to interact with the A subunit with high specificity; indeed, this fragment was used for the affinity purification of V-ATPases from cell and tissue extracts (Fig. 1A; see also Abbas et al., 2020). GFP or mCherry were linked to SidK_1-278_ via its C terminus because the structure of the V-ATPase:SidK_1-278_ complex suggested that attachment at this position was unlikely to affect association of the chimera with the V-ATPase (Fig. 1B; Abbas et al., 2020). For brevity, the resulting construct is referred to simply as SidK.

**Figure 1.**
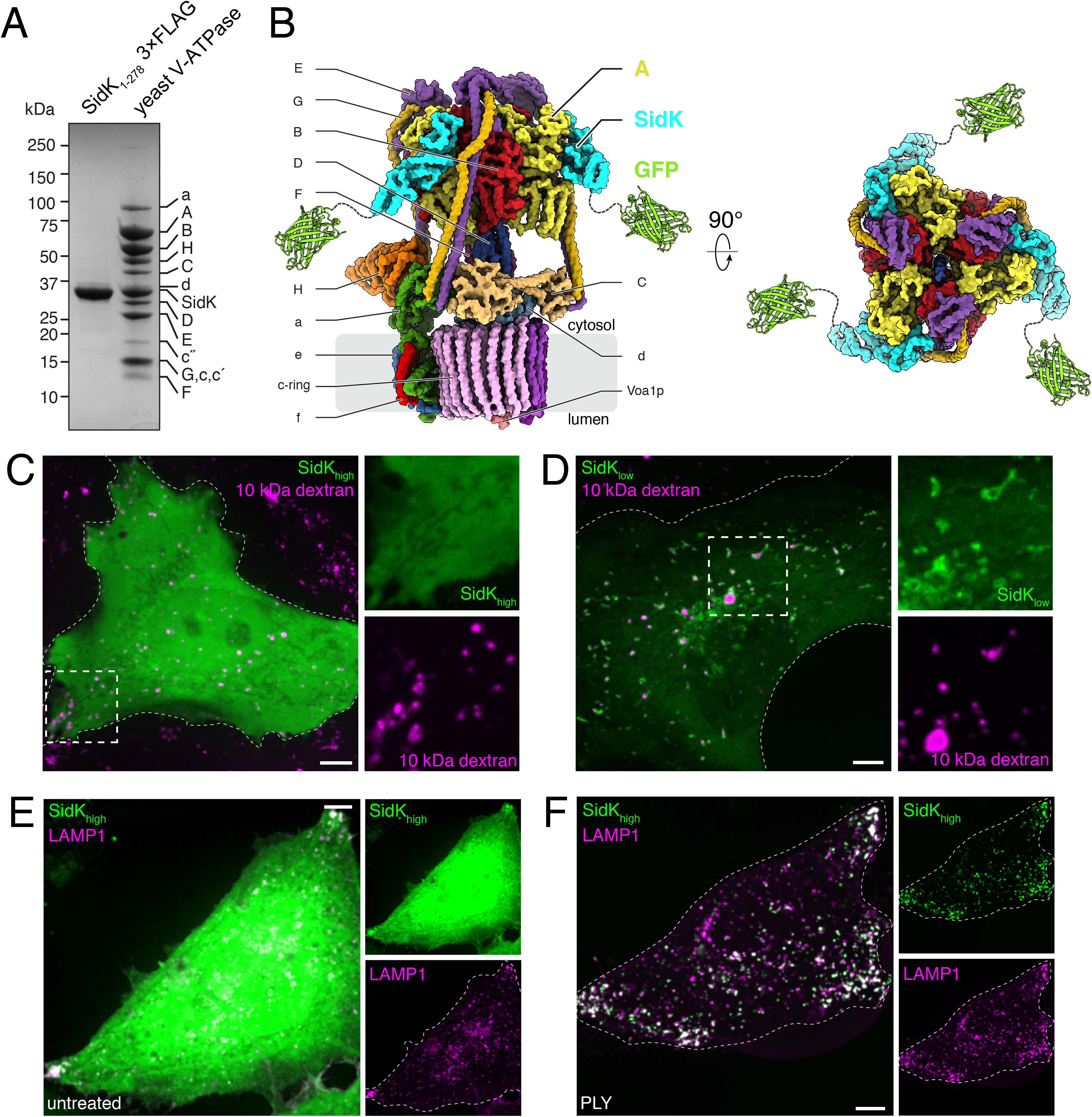
SidK interacts with the V-ATPase *in vitro* and when expressed in mammalian cells. **A.** SDS-PAGE of 3×FLAG SidK_1-278_ to illustrate purity of the probe and its ability to allow isolation of yeast V-ATPase by affinity chromatography. Bands corresponding to SidK_1-278_ and to individual subunits of V-ATPase are labeled. **B.** Model of the yeast V-ATPase with SidK_1-278_-GFP bound, generated from PDB accession numbers 5VOX, 1GFL, and 6O7T. **C.** Visualization of SidK_high_ cells. Lysosomes of HeLa cells were transfected with SidK-GFP (green) and loaded with Alexa Fluor-647-labelled 10 kDa dextran (magenta). Side panels show the individual SidK and 10 kDa dextran channels in the region denoted by the dotted square, at 2.2× magnification. **D.** Visualization of SidK_low_ cells (green) labeled for lysosomes as in **C**. Side panels show the individual SidK and 10 kDa dextran channels denoted by the dotted square, at 2.0× magnification. **E** and **F.** Lysosomes were identified by LAMP1 immunostain (magenta) after expression of high levels of SidK-GFP (SidK_high_; green). Prior to fixation and immunostaining, cells were either left untreated (**E**) or permeabilized with pneumolysin (PLY) to remove excess cytosolic SidK-GFP (**F**). Side panels show the individual SidK and LAMP1 channels. Here and elsewhere, outlines of cells are indicated by dotted lines when required. Images in **C-F** are extended focus compressions of confocal images representative of ≥ 30 fields from ≥ 3 separate experiments of each type. All scale bars: 5 μm.

As is often the case, varying levels of expression were observed following transient transfection of fluorescent SidK in HeLa cells. The fluorescence intensity of the soluble (excess) SidK seen in high expressers (SidK_high_) precluded the resolution of the fraction of the construct bound to V-ATPase-containing organelles, such as lysosomes (Fig. 1C). In contrast, SidK showed clear association with vesicular and tubular structures in low-expressing cells (SidK_low_, Fig. 1D), some of which were identifiable as lysosomes by loading with fluorescent dextran. Because the excess fluorescence of SidK_high_ cells appeared to be cytosolic, we predicted that selective removal of soluble material would reveal the more tightly-bound, organelle-associated probe. This assumption was tested by comparing SidK_high_ cells before (Fig. 1E) and after (Fig. 1F) selective leaching of cytosolic components following permeabilization of the plasma membrane with the pore-forming toxin, pneumolysin (PLY). As anticipated, the cytosolic fluorescence was largely depleted in PLY-treated cells, revealing membrane-associated SidK, a fraction of which was clearly co-localized with LAMP1, a marker of late endosomes/lysosomes. We concluded that the fluorescent chimeras of SidK can be used for detection of membrane-associated ligands, presumably V-ATPases, particularly in cells with low expression level or permeabilized with PLY or similar reagents.

### Expression of SidK-fluorescent protein constructs affects lysosomal positioning and pH

In otherwise untreated HeLa cells late endosomes/lysosomes are largely located near the nucleus, as revealed by LAMP1 immunostaining (Fig. 2A). However, we noted that endo/lysosomes tended to accumulate at the cell periphery of SidK-transfected cells, particularly those where the construct was highly expressed (Fig. 2B; see also 1F). Because SidK has been shown to inhibit the V-ATPase partially both *in vitro* and in mammalian cells (Zhao et al., 2017; Xu et al., 2010), we considered the possibility that inhibition of proton pumping was responsible for the altered distribution of LAMP1-positive compartments. This notion was validated by treating cells with concanamycin A, a potent and specific V-ATPase inhibitor, which resulted in a similar margination of a fraction of the LAMP1-positive compartments (Fig. 2C). These observations are consistent with previous reports that the positioning of lysosomes correlates with their luminal pH (Johnson et al., 2016).

**Figure 2.**
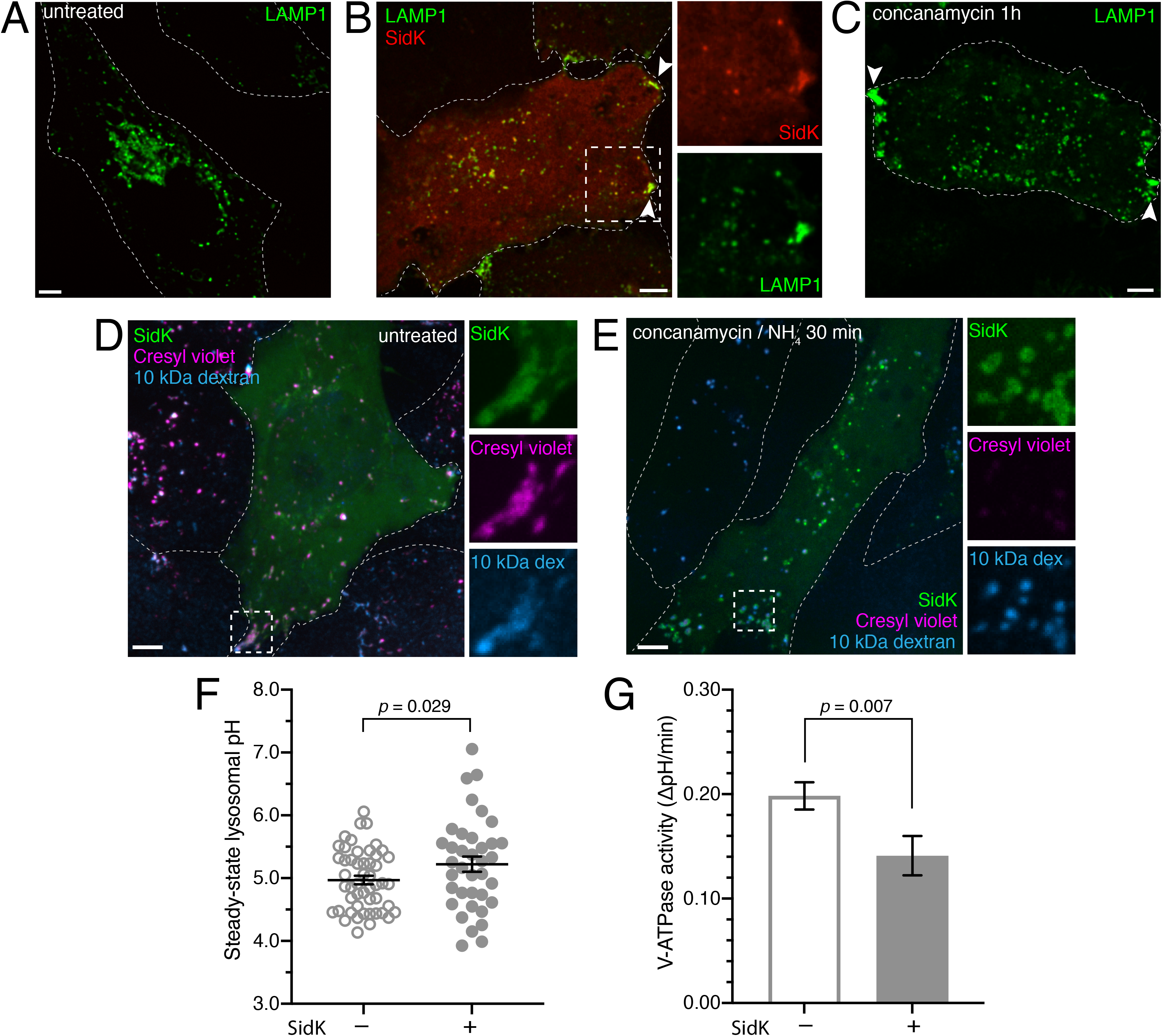
Effect of SidK-fluorescent protein overexpression on the pH of acidic compartments. **A.** Localization of endogenous LAMP1 (green) in untreated control cells. **B.** Localization of LAMP1 (green) in cells expressing SidK-mCherry (red). Side panels show the individual SidK and LAMP1 channels in the area denoted by the dotted square, at 2.0× magnification. Arrowheads mark sites of peripheral LAMP1 accumulation. **C.** Localization of LAMP1 (green) in cells treated with 250 nM concanamycin A for 1 hr. Arrowheads mark sites of peripheral LAMP1 accumulation. **D.** Retention of the acidotropic dye cresyl violet (magenta) in cells expressing SidK-GFP (green), where lysosomes had been preloaded with Alexa Fluor-647-conjugated 10 kDa dextran (blue). Side panels show the individual channels of the area denoted by the dotted square, at 2.7× magnification. **E.** Cells expressing SidK-GFP (green) and labeled for lysosomes (blue) were treated with 250 nM concanamycin A and 10 mM NH_4_Cl for 30 min to neutralize luminal pH. After treatment, cells were incubated with cresyl violet as in **D**. Side panels are 2.7× magnification. Images in **A-E** are extended focus images representative of ≥ 30 fields from ≥ 3 separate experiments of each type. All scale bars: 5 μm. **F.** Lysosomal pH was determined in SidK^−^ and SidK^+^ HeLa cells. The lysosomes of control or SidK-mCherry-transfected HeLa cells were loaded overnight with FITC-10 kDa dextran. Lysosomal pH was subsequently measured by ratiometric fluorescence microscopy as described in the Materials and Methods. For each condition, 3 independent experiments were quantified, with ≥ 20 cells per replicate. Data are means ± SEM. *p* value was calculated using unpaired, 2-tailed student’s t-test. **G.** Lysosomal V-ATPase activity was measured in control and SidK-mCherry-expressing HeLa cells acutely treated with 500 nM concanamycin A. V-ATPase activity was determined from the inverse rate of alkalization upon the addition of concanamycin A. See Materials and Methods for further details. For each condition, three independent experiments were quantified, with ≥ 10 cells per replicate. Data are means ± SEM. *p* value was calculated using unpaired, 2-tailed student’s t-test.

To verify that SidK caused lysosomal alkalinization we used cresyl violet (Ostrowski et al., 2016), an acidotropic fluorescent dye that is more photostable than the LysoTracker dye used by Xu et al. (2010). Unexpectedly, and in apparent disagreement with the findings of Xu et al. (2010), the lysosomes remained acidic (retained cresyl violet) despite the presence of SidK in the cytosol (Fig. 2D). The reliability of cresyl violet as an indicator of lysosomal acidification was confirmed using concanamycin A, which prevented the accumulation of the acidotropic dye (Fig. 2E). It is noteworthy, however, that like other acidotropic dyes, cresyl violet is a coarse indicator of pH, unable to sense moderate changes in pH. In this regard, it is of interest that *in vitro* determinations using purified yeast V-ATPase showed that saturating concentrations of SidK caused only a ≈ 30% inhibition of ATPase activity (Zhao et al., 2017). To more precisely assess the effect of SidK expression on HeLa cells, their lysosomes were loaded with FITC-dextran and pH was measured ratiometrically *in situ* (see Materials and Methods). Using this approach, the pH of lysosomes in SidK-expressing cells was found to be moderately –yet significantly– more alkaline (pH 5.22 ± 0.12) than that of control cells (pH 4.97 ± 0.07; Fig. 2F). As discussed above, a variety of parameters influence the steady-state luminal pH of an organelle. Therefore, we endeavored to more directly assess the effect of SidK on proton pumping by the V-ATPase. This activity can be measured by quantifying the effect of concanamycin on proton flux across the lysosomal membrane. These measurements are based on the assumption that at steady-state pH, V-ATPase activity is precisely offset by an equivalent but opposite backflux (leak) of protons. The initial rate of proton leakage unmasked by addition of saturating concentrations of concanamycin can therefore be considered as an accurate measure of proton pumping by the V-ATPase at steady-state. This analysis (Fig. 2G) indicated that SidK reduced proton pumping by the V-ATPase by 29%, in good agreement with previous *in vitro* measurements (Zhao et al., 2017). Whether the modest change in pH or the more obvious inhibition of the rate of pumping are responsible for the redistribution of lysosomes within the cells remains to be established.

### Localization of the V-ATPase in mammalian cells using SidK-AL568

The preceding results indicated that, while capable of detecting the V-ATPase under appropriate conditions, expression of a genetically-encoded form of fluorescent SidK had obvious limitations, most notably the fact that chronic inhibition of the pump –even if only partial– alters the distribution of the labeled organelles. To avoid inhibition of the pump prior to its detection we developed an alternative probe based on SidK that could be used to label the V-ATPase in fixed and permeabilized cells. Recombinant SidK_1-278_ was expressed in bacteria and, after purification to near homogeneity (Fig. 1A), was covalently labeled with Alexa Fluor 568 (referred to hereafter as SidK-AL568) and used to stain cells. In HeLa cells, SidK-AL568 labelled vesicular and cisternal compartments reminiscent of endosomes and the Golgi complex with remarkably little background noise (Fig. 3A, left panel). This staining was blocked by pretreatment with unlabeled SidK (Fig. 3A, right panel), implying that binding is saturable and that Alexa Fluor conjugation did not alter the binding properties of SidK.

**Figure 3.**
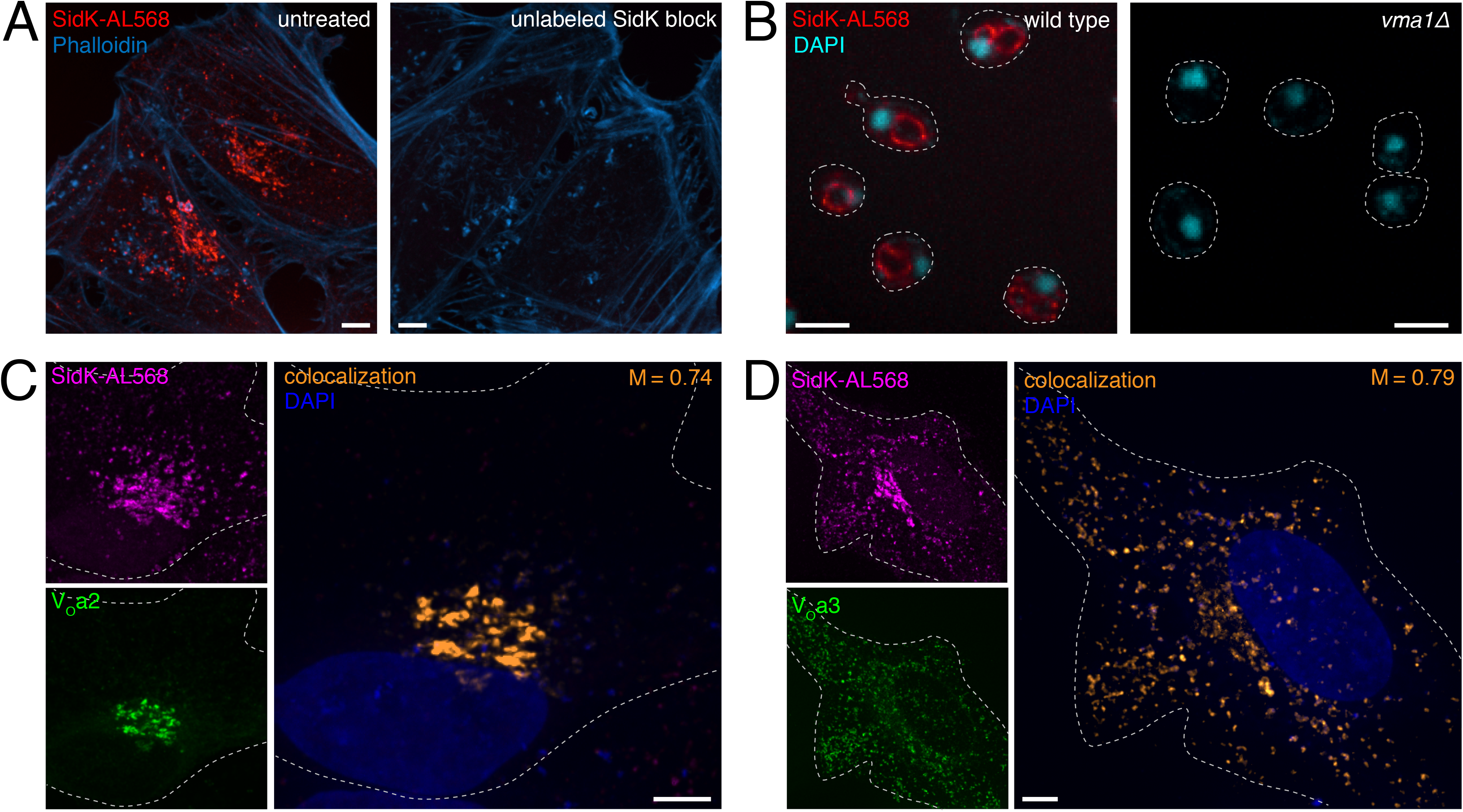
SidK-AL568 specifically labels the V-ATPase. **A.** HeLa cells were stained with SidK-AL568 (red) as described in the Materials and Methods either directly (left) or after incubation with a 5-fold excess of unlabeled SidK_1-278_ (right). F-actin was labeled with fluorescent phalloidin (blue). **B.** Spheroplasts of wild type or *vma1*Δ *S. cerevisiae* were prepared and stained with SidK-AL568 (red) as detailed in Materials and Methods. Nuclei were labeled with DAPI (cyan). Outlines of the spheroplasts, visualized by DIC, are indicated by dotted lines. **C** and **D.** HeLa cells were transfected with V_O_a2- (**C**) or V_O_a3-GFP (**D**) constructs (green), and stained using SidK-AL568 (magenta). Nuclei were labeled with DAPI (blue). Outlines of cells are indicated by dotted lines. Side panels show the individual SidK-AL568 and V_O_a channels. Colocalization between SidK-AL568 and the corresponding V_O_a subunit is shown in orange and the calculated Manders’ coefficient (M) between the a subunit and SidK-AL568 is indicated. **A-D** are extended focus compressions of confocal images representative of ≥ 30 fields from ≥ 3 separate experiments of each type. All scale bars: 5 μm.

The specificity of SidK-AL568 for the A subunit of the V-ATPase was validated by comparing staining of wild-type *Saccharomyces cerevisiae* to that of a mutant strain lacking the gene for subunit A (*vma1*Δ). SidK-AL568 clearly labeled the vacuole and pre-vacuolar compartment of wild-type spheroplasts, but staining was absent in the *vma1*Δ strain. Additional evidence that SidK-AL568 associates with assembled V-ATPase complexes in mammalian cells was obtained by ectopic (over)expression in HeLa cells with fluorescently tagged V_o_a2 or V_o_a3 subunits that localize predominantly to the Golgi complex or endosomes, respectively (Saw et al., 2011). SidK-AL568 co-localized with both V_o_a2- and V_o_a3-GFP, with highly significant Manders’ coefficients (M= 0.74 and 0.79, respectively; Figs. 3C and D). Taken together, these data verified that fluorescently labeled SidK is a sensitive tool for the specific detection of the V-ATPase in eukaryotic cells, where it detects primarily organelle-associated complexes.

We proceeded to use SidK-AL568 to assess the presence and density of V-ATPase complexes in defined intracellular compartments, an experiment that –while conceptually simple– has been hampered by the paucity of sufficiently sensitive reagents. As intimated above and illustrated in more detail in Fig. 4A, SidK-AL568 stained peripheral vesicular compartments (Fig. 4A, open arrowheads) as well as juxtanuclear vesicles and cisternae (closed arrowheads) likely corresponding to endocytic and Golgi components, respectively. These assumptions were confirmed by simultaneously visualizing LAMP1 and the *trans*-Golgi (Figs. 4B and C), demonstrating that SidK-AL568 staining has a high degree of colocalization with these compartments, which are known to have a markedly acidic lumen (Casey et al., 2009). In contrast, SidK-AL568 showed minimal colocalization with markers of the endoplasmic reticulum (ER) and mitochondria (Figs. 4E and F), which have a near-neutral or slightly alkaline lumen (Casey et al., 2009). Of note, SidK-AL568 labeled poorly the *cis*-Golgi (Fig. 4D), which is thought to be less acidic than the *mid*- and *trans*-cisternae.

**Figure 4.**
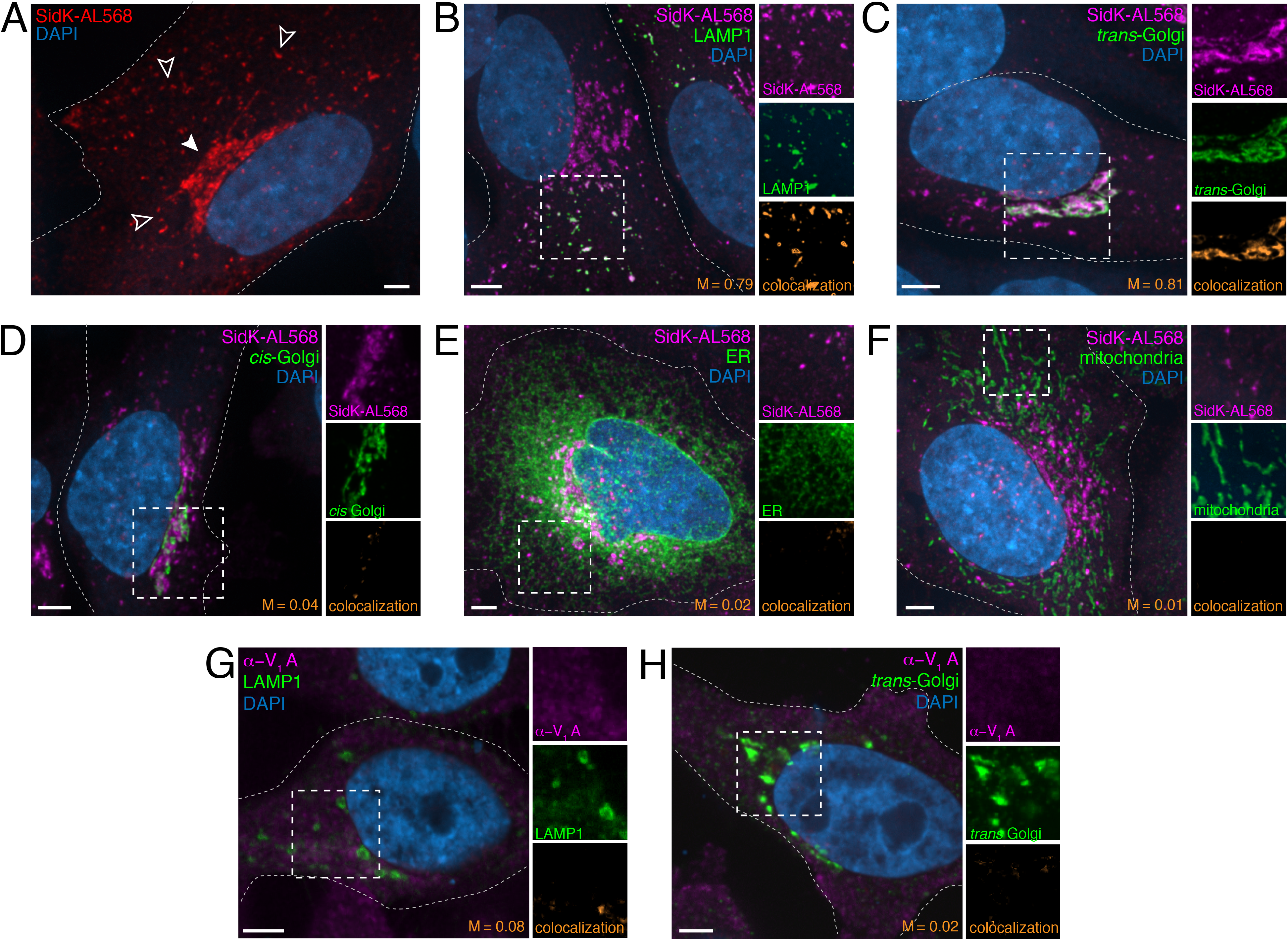
Localization of the V-ATPase in mammalian cells using SidK-AL568. **A**. HeLa cells were stained with SidK-AL568 (red), as described in Materials and Methods. Nuclei were labeled with DAPI (blue). Arrowheads mark peripheral, likely endocytic, vesicles (open) and juxtanuclear vesicles and cisternae, likely including the Golgi compartment (closed). **B-F.** V-ATPase in HeLa cells was labeled with SidK-AL568 (magenta) and co-stained for **B**: LAMP1, **C:** *trans*-Golgi, **D:** *cis*-Golgi, **E:** ER, and **F:** mitochondria (green) as described in the text. Nuclei were labeled with DAPI (blue). Side panels show the individual SidK-AL568 and organelle channels in the area denoted by the dashed square, at 1.1, 0.9, 1.1, 1.7 and 1.5× magnification, respectively. Outlines of cells are indicated by dotted lines. **G** and **H.** HeLa cells were immunostained using a commercially available V_1_A antibody (Abnova; catalogue no. H00000523-M02; shown in magenta) in cells co-stained for LAMP1 (**G**) or *trans*-Golgi (**H**), in green. Nuclei were labeled with DAPI (blue). Side panels show the individual α-V_1_A and organelle channels marked by the dotted square, at 1.1 and 1.4× magnification, respectively. **A-H** are extended focus images representative of ≥ 30 fields from ≥ 3 separate experiments of each type. Outlines of cells are indicated by dotted lines. Colocalization and Manders’ coefficients (M) between the organellar marker and V-ATPase probe are shown in orange. All scale bars: 5 μm.

Based on the previous observations, we believe that SidK-AL568 is an excellent probe to detect V-ATPase, possibly superior to other reagents commonly used in the literature. Indeed, when compared to a commercially available antibody raised against the same subunit to which SidK-AL568 binds (V-ATPase subunit A), our probe yielded better results. The antibody chosen for comparison (α-ATP6V1A) is widely used in the literature to document the localization of V-ATPases, often as the basis to reach important functional conclusions (e.g. Ramirez et al., 2019). When tested in HeLa cells (Figs. 4G and H), this antibody yielded a diffuse punctate pattern reminiscent of that reported by Ramirez et al. (2019) for tumour cells and elsewhere for a variety of other cells stained with other A subunit antibodies (Yajima et al., 2007; McGuire et al., 2019; Michel et al., 2013). However, its co-localization with acidic compartments was poor: the Manders’ coefficient of the antibody with LAMP1 was M = 0.08 (compared to an M = 0.79 with SidK-AL568), while that with anti-TGN46, the *trans*-Golgi marker used, was M = 0.05 (compared to an M = 0.81 with SidK-AL568). Comparison of the *α*-ATP6V1A signal with that of SidK-AL568 showed that most of the α-ATP6V1A staining was background, which was removed by applying the unbiased Costes thresholding method prior to colocalization analysis (Costes et al., 2004); see Materials and Methods); this was not the case for SidK-AL568 staining (Fig. S1). We therefore suggest that SidK-AL568 is a more specific, preferable probe.

### Assessment of V-ATPase acquisition by maturing phagosomes

The phagosomes formed by cells of the innate immune system, such as macrophages, are specialized compartments with microbicidal and degradative functions. The lumen of nascent phagosomes is near-neutral, but becomes gradually acidic as the compartment matures, reaching a pH ≤ 5 (e.g. Fig. 5A). This acidification has been demonstrated to depend on the activity of V-ATPases (Lukacs et al., 1990), which are nevertheless undetectable on the macrophage plasma membrane that forms the initial phagosomal enclosure. The graded acidification is thought to result from accumulation of V-ATPases complexes owing to fusion of the nascent phagosome with early and late endosomes and, ultimately, with lysosomes. Remarkably, to our knowledge, this purported mechanism has not been documented experimentally. We therefore utilized the SidK-AL568 probe to detect V-ATPases during the phagosomal maturation process, using murine macrophages (RAW264.7 cells) that had been transfected with various membrane markers correlated to maturation state (Fig. 5B).

**Figure 5.**
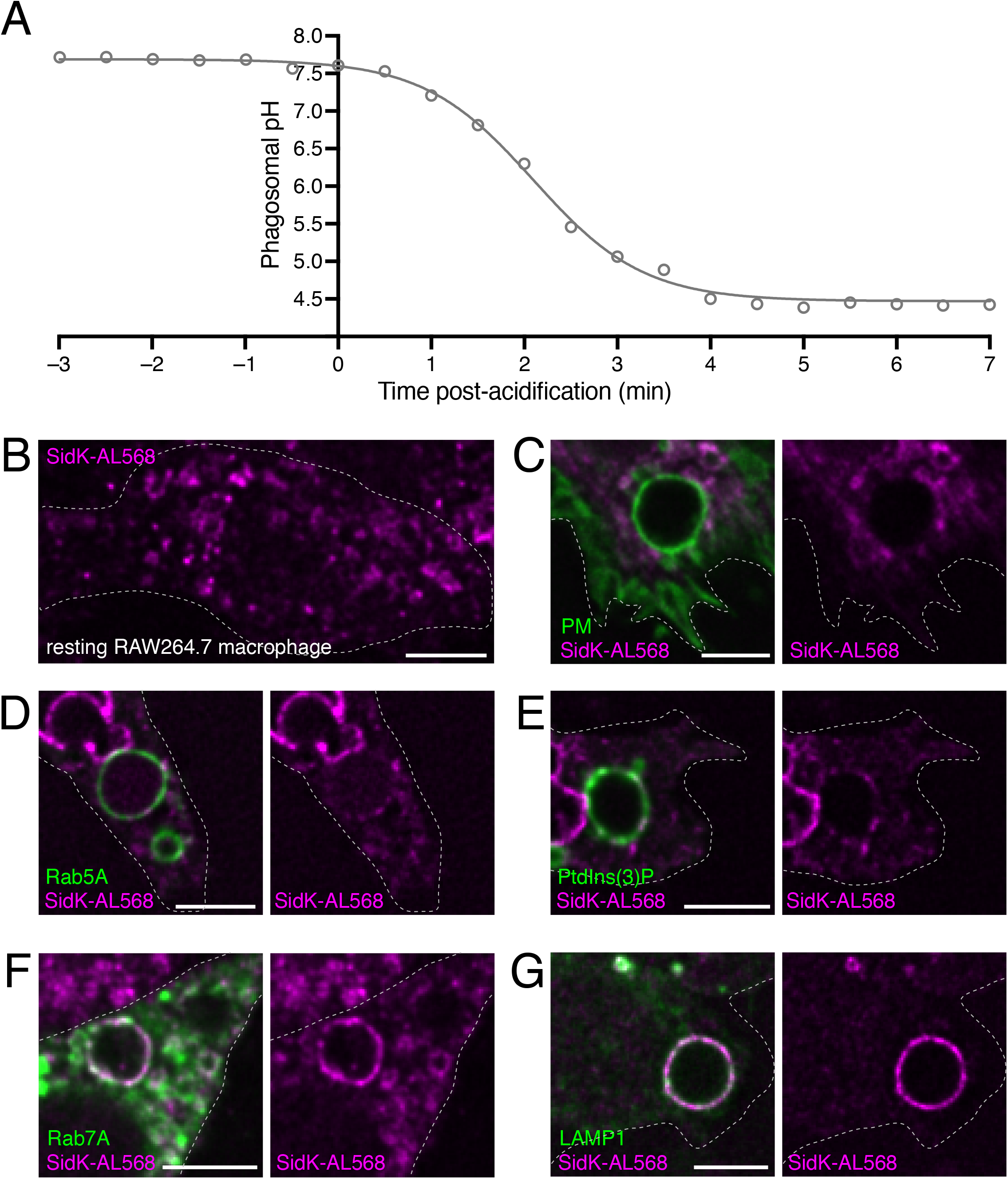
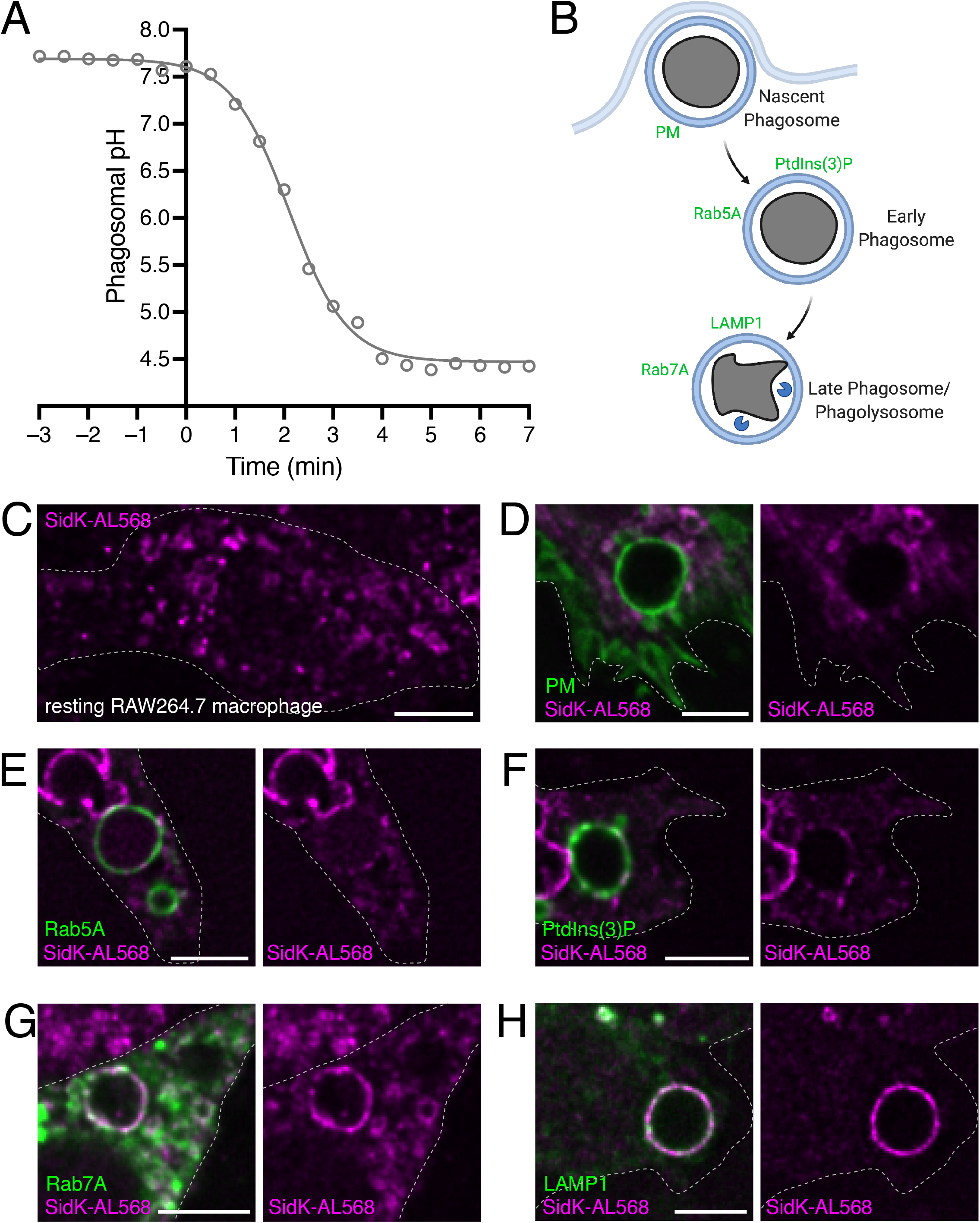
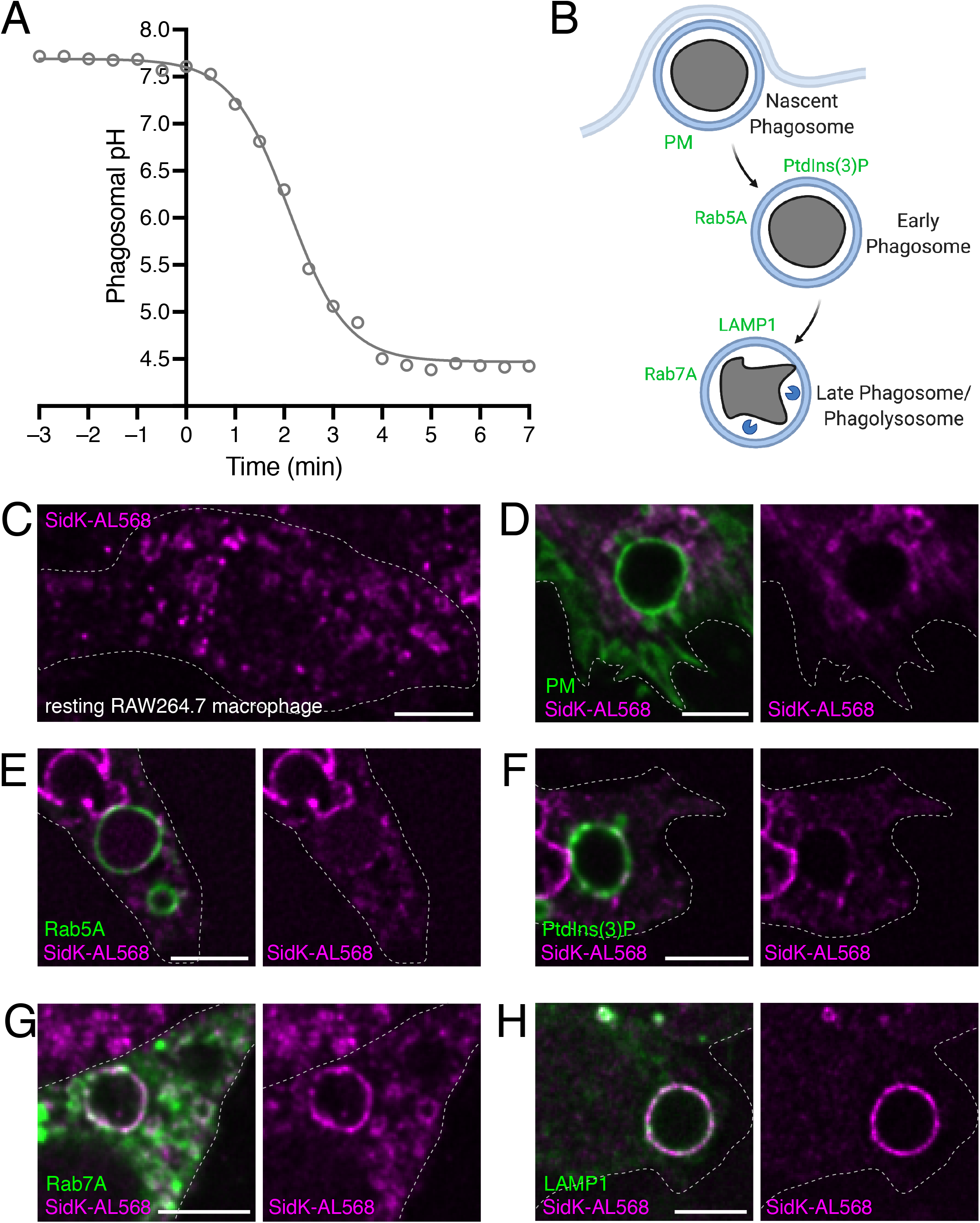
The acidification of macrophage phagosomes parallels the recruitment of the V-ATPase to the phagosomal membrane, as detected by SidK-AL568. **A.** RAW264.7 cells were incubated with IgG-opsonized FITC-zymosan, which were sedimented by centrifugation onto the coverslip to initiate phagocytosis. Phagosomal pH changes were determined by ratiometric fluorescence microscopy, as described in the Materials and Methods. Time 0 marks the point where phagosome acidification became evident. Data points (open circles) represent the average of 2 independent experiments, with ≥ 10 cells per replicate. **B.** Schematic showing the stages of phagosome engulfment and maturation, with associated membrane markers. **C.**V-ATPase localization in resting RAW264.7 cells stained with SidK-AL568 (magenta). **D-H**. RAW264.7 cells that had been transfected with the indicated phagosomal maturation markers (green) were allowed to internalize IgG-opsonized SRBCs for various times before fixation. The cells were then fixed, permeabilized, and stained with SidK-AL568 (magenta). **D.** Cells expressing a plasma membrane (PM) marker, fixed 2 min after initiation of phagocytosis. **E.** Cells expressing Rab5A, fixed 5 min after initiation of phagocytosis. **F.** Cells expressing a PtdIns(3)P-specific probe, fixed 5 min after initiation of phagocytosis. **G.** Cells expressing Rab7 marker, fixed 30 min after initiation of phagocytosis. **H.** Cells immunostained for LAMP1, fixed 30 min after initiation of phagocytosis. **C-H** Are *XY* optical slices acquired near the middle of the cell or phagosome, representative of ≥ 30 fields from ≥ 3 separate experiments of each type. Outlines of cells are indicated by dotted lines. All scale bars: 5 μm.

Like HeLa cells, resting RAW264.7 macrophages showed distinct vesicular staining with SidK-AL568, as well as larger vacuoles that likely form by macropinocytosis, which is constitutively active in these cells (Fig. 5C). Nascent phagosomes, enriched by arresting phagocytosis shortly (2 min) after exposure to the target particles, were identified by the persistence of plasmalemmal markers (e.g. PM-GFP). As expected, these nascent phagosomes were essentially devoid of V-ATPases, although V-ATPase-rich organelles seemed to accumulate in their immediate vicinity (Fig. 5D). In contrast, early phagosomes –which are measurably acidic and were identified by the acquisition of Rab5 and PtdIns(3)P– showed distinct acquisition of SidK-AL568 into discrete areas (Fig. 5E and F). Late phagosomes/phagolysosomes, which were identified by possessing Rab7 or LAMP1, were even more enriched with SidK-AL568 (Fig. 5G and H), consistent with their highly acidic pH. Interestingly, a patchy localization of SidK-AL568 was noted at all stages of phagosome maturation, suggesting that the recruitment of V-ATPases may occur at restricted sites, which may have important implications for traffic and pH regulation.

### The V-ATPase is heterogeneously distributed on the membrane of acidic organelles

The discontinuous pattern of SidK-AL568 on the late-phagosome membrane is reminiscent of the spatial segregation of other membrane components that was observed previously and associated with the formation of ER-phagosome contacts (Levin-Konigsberg et al., 2019). These contacts are established, at least partly, by interaction of phagosomal ORP1L with the ER resident proteins VapA and VapB (Rocha et al., 2009; Loewen and Levine, 2005). We considered the possibility that the SidK-AL568 patches represented similar regions of V-ATPase exclusion from ER contact sites. To assess this possibility, phagosomes of RAW264.7 cells that had been transfected with ORP1L-GFP were stained with SidK-AL568. Following transfection (Fig. 6A), ORP1L and the V-ATPase showed an inverse distribution, the latter accumulating in regions where the former was depleted. This segregation could be quantified and is represented as a ratio of SidK-568:ORP1L fluorescence in the rightmost panel of Fig. 6A. In contrast, SidK-AL568 co-distributed with Arl8b (Fig. 6B), which was shown earlier to be excluded from ER contact sites (Levin-Konigsberg et al., 2019). It was imperative to ensure that the apparent depletion of V-ATPases from contact areas was not caused by limited access of SidK-AL568. Accessibility was verified by immunostaining VapB alongside SidK-AL568 (Fig. 6C). As was the case for ORP1L, the regions that were rich in VapB were comparatively depleted of SidK-AL568. Because the antibody used to immunostain VapB is much larger (≈150 kDa) than SidK-AL568 (≈35 kDa), exclusion of the V-ATPase ligand cannot account for the observed segregation, which is an indication of the genuine existence of V-ATPase-enriched microdomains.

**Figure 6.**
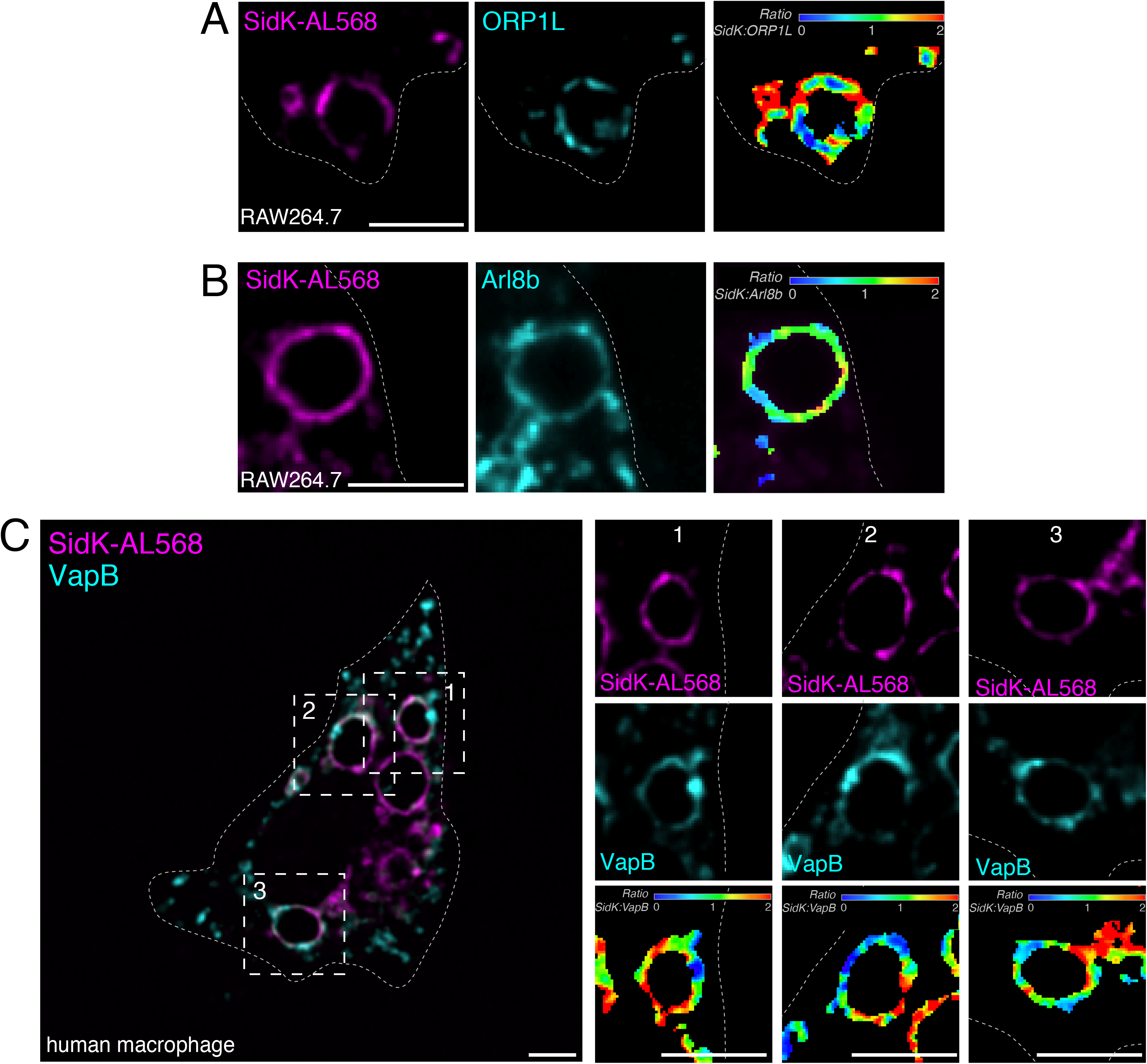
V-ATPase accumulates in subdomains along the phagosomal membrane and is excluded from ER contact sites. **A.** RAW264.7 cells that had been transfected with ORP1L-GFP were allowed to internalize IgG-opsonized SRBCs and fixed after 60 min. Cells were stained with SidK-AL568 (magenta) and visualized along with ORP1L (cyan). Individual SidK-AL568 and ORP1L channels and the SidK-AL568:ORP1L fluorescence ratio (*Ratio_SidK- AL568:ORP1L_*) are shown from left to right. Fluorescence ratio is pseudocolored in a rainbow LUT, corresponding to ratio values from 0 to 2. **B.** RAW264.7 cells that had been transfected with Arl8b-GFP were allowed to internalize IgG-opsonized SRBCs and fixed after 30 min. Cells were stained with SidK-AL568 (magenta) and visualized along with Arl8b-GFP (cyan). Individual channels and the pseudocolored fluorescence ratio are shown from left to right. **C.** Human M0 macrophages were allowed to internalize IgG-opsonized SRBCs and fixed after 30 min. Cells were fixed, permeabilized and stained with SidK-AL568 (magenta) and immunostained for endogenous VapB (cyan). Smaller panels show individual SidK-AL568, VapB and SidK-AL568:VapB fluorescence ratio channels (*Ratio_SidK-AL568:VapB_*) (top to bottom) for 3 individual phagosomes (left to right) identified by the dotted squares, all at 1.8× magnification. **A-C** are central *XY* optical slices optical slices acquired near the middle of the cell or phagosome representative of ≥ 30 fields from ≥ 3 separate experiments of each type. Outlines of cells are indicated by dotted lines. All scale bars: 5 μm.

The realization that unappreciated microdomains rich in V-ATPases exist in phagosomes prompted us to ask whether similar subdomains exist in other organelles. To facilitate visualization, we initially generated enlarged lysosomes using sucrose (Cohn and Ehrenreich, 1969; Bright et al., 1997; DeCourcy and Storrie, 1991; Swanson et al., 1986; Ferris et al., 1987) in cells that had been transfected with ORPL1 (Fig. 7A) or Arl8b (Fig. 7B). As with late phagosomes, V-ATPase (SidK-AL568) was depleted from membrane domains where ORPL1 was found, while coinciding with Arl8b.

**Figure 7.**
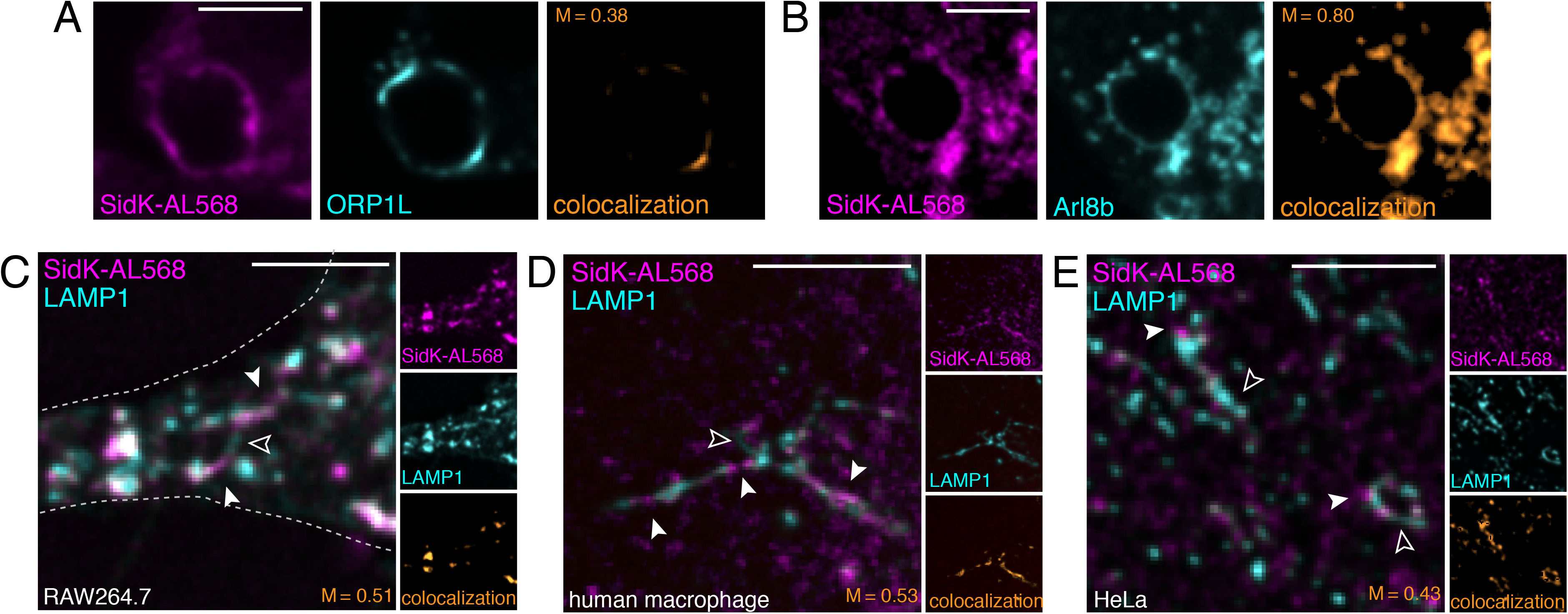
Lysosomes and lysosomal tubules in a variety of cell types show V-ATPase subdomains. **A.** RAW264.7 cells transfected with ORP1L-GFP were subjected to lysosome enlargement by overnight treatment with 30 mM sucrose, as described in the Materials and Methods. Cells were stained with SidK-AL568 (magenta) and the distribution of V-ATPase in sucrosomes labeled with ORP1L (cyan) visualized. Side panels show the individual SidK-AL568, ORP1L and colocalization channels (left to right). Colocalization and Manders’ coefficient between ORP1L and SidK-AL568 (M) are shown in orange. **B.** RAW264.7 cells transfected with Arl8b-GFP were subjected to lysosome enlargement by overnight treatment with 30 mM sucrose. Cells were stained with SidK-AL568 (magenta) and the distribution of V-ATPase in sucrosomes labeled with Arl8b (cyan) visualized. Side panels show the individual SidK-AL568, Arl8b and colocalization channels (left to right). Colocalization and Manders’ coefficients between Arl8b and SidK-AL568 (M) are shown in orange. **C-E** Localization of the V-ATPase in tubular lysosomal structures in: **C.** RAW264.7 cells, **D.** human M0 macrophages, and **E.** HeLa cells. Cells were stained using SidK-AL568 (magenta) and immunostained for endogenous LAMP1 (cyan). Solid white arrowheads mark lysosomal regions that are SidK^+^ LAMP1^−^, while open arrowheads mark regions that are SidK^−^ LAMP1^+^. Side panels show individual SidK-AL568, LAMP1 and colocalization channels (top to bottom). Colocalization and Manders’ coefficients between LAMP1 and SidK-AL568 (M) are shown in orange. **A-E** are *XY* optical slices representative of ≥ 30 fields from ≥ 3 separate experiments of each type. All scale bars: 5 μm.

In addition to visualizing V-ATPase microdomains in enlarged phagosome or sucrose-enlarged lysosomes, the existence of comparatively long tubular lysosomes in macrophages also enabled us to assess segregation in resting, unmodified cells. In RAW264.7 macrophages (Fig. 7C) and human monocyte-derived macrophages (Fig. 7D), regions of late endosome/lysosomes –identified by LAMP1– were preferentially enriched with V-ATPase. This pattern could also be observed in some HeLa cells where the LAMP1-positive structures were sufficiently large (Fig. 7E). In all cases, regions of V-ATPase exclusion from LAMP1-stained areas were observed; in some instances SidK-AL568-stained structures appeared to bud off from LAMP-positive tubular lysosomes. The observed segregation may be an indication of selective delivery or removal of V-ATPases at varying stages of organellar maturation.

### Quantification of the number of V-ATPases per lysosome: evidence of heterogeneous density that correlates with the subcellular localization of the organelles

To date, estimates of the number of V-ATPase complexes in vesicular compartments have been global approximations based on measurements made in pooled whole-cell preparations (de Araujo et al., 2020; Takamori et al., 2006). In an effort to refine estimates and provide topological information, we attempted to quantify the number of SidK-AL568 molecules bound per organelle. This analysis required determination of the fraction of SidK molecules labeled by the Alexa dye and the number of fluorophores attached per SidK-AL568 molecule, followed by comparison of the single molecule fluorescence to the total fluorescence associated with the organelle of interest. We determined that 96.7% of the molecules were labeled and analysis of the photobleaching pattern of monodisperse SidK (Fig. 8A; see Materials and Methods) indicated that 99% of these molecules had reacted with a single fluorophore (Fig. 8B).

**Figure 8.**
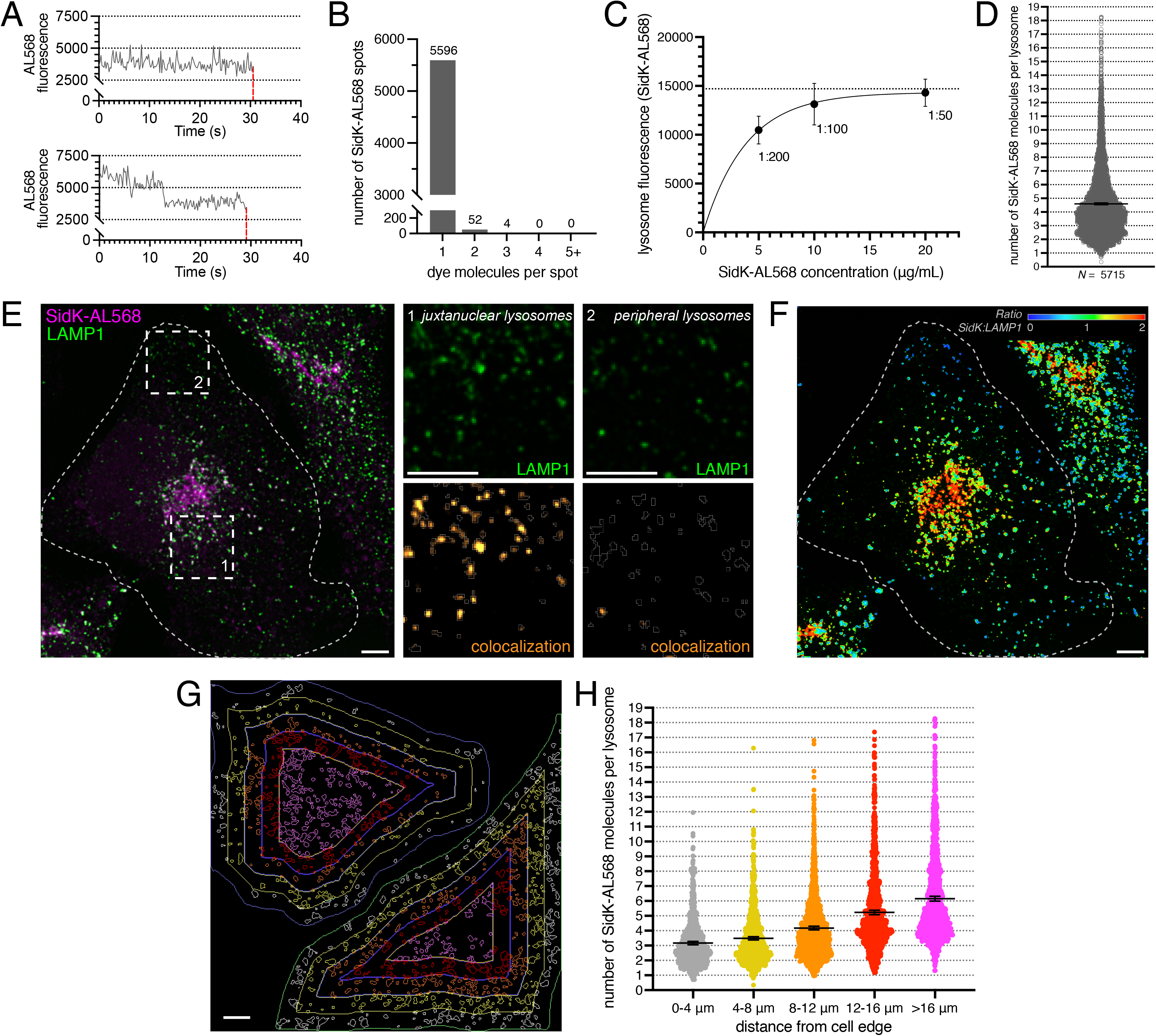
Estimation of the number of V-ATPases per lysosomal compartment. **A.** Photobleaching of monodisperse SidK-AL568. Representative intensity-time traces for a single SidK-AL568 labeled with 1 (top) or 2 (bottom) Alexa Fluor 568 moieties. Loss of fluorescence after complete photobleaching is shown by the dotted red lines. **B.** Histogram showing the measured number of dye molecules per SidK-AL568 monomer determined analyzing > 5000 single molecule photobleaching traces. The number of SidK-AL568 molecules in each category of the histogram is indicated above the bar. **C.** HeLa cells were stained with SidK-AL568 at varying concentrations (shown in absolute concentration and as dilution). Lysosomal fluorescence was plotted to determine the concentration of SidK-AL568 required to reach maximal lysosome fluorescence, indicative of saturation of available binding sites. **D.** HeLa cells were stained with SidK-AL568 using the near-saturation conditions determined in **C** (1:50). Total fluorescence per lysosome was compared to the average fluorescence of a single molecule of SidK-AL568 and is displayed; the average number of SidK-AL568 molecules bound per lysosome is indicated by the horizontal line, as is the SE (whiskers). A total of 5715 lysosomes from 2 independent experiments were quantified. **E.** Intracellular V-ATPase distribution among lysosomes is heterogeneous. HeLa cells were stained with SidK-AL568 (magenta) using the near-saturating conditions determined in **C** (1:50), and co-stained for LAMP1 (green). Side panels show the individual LAMP1 (top), and SidK-AL568:LAMP1 colocalization channels (bottom) for juxtanuclear (left) or peripheral (right) lysosomes in the areas denoted by the dotted squares. Outlines of cells and lysosomes are indicated by dotted or solid lines, respectively. **F.** SidK-AL568:LAMP1 fluorescence ratio (*Ratio_SidK-AL568:LAMP1_*) corresponding to image in **E**. The fluorescence ratio was pseudocolored in a rainbow LUT, representing ratio values from 0-2. **G.** Shell analysis of SidK distribution as a function of distance from the cell outer edge. HeLa cells were stained using SidK-AL568 and co-stained for LAMP1. Cell outlines were drawn and degraded iteratively inward by 4 μm to create concentric shells (colored differentially) within each cell that were used to subgroup LAMP1+ lysosomes for subsequent analysis. **H.** For each shell, total fluorescence per lysosome was compared to the average fluorescence of a single molecule of SidK-AL568, and the average number of SidK-AL568 molecules bound per lysosome calculated, as a function of distance from the cell outer edge. Data are means ± SEM of 2 independent experiments with ≥ 5 cells per replicate. *N* = 791, 1112, 1203, 1275 and 1334 lysosomes analyzed for shells 0-4, 4-8, 8-12, 12-16, and >16 μm from the cell edge, respectively. Images in **E-G** are extended focus compressions of confocal images representative of ≥ 10 fields from ≥ 2 separate experiments of each type. All scale bars: 5 μm.

We next determined the concentration of SidK-AL568 needed to saturate all the available binding sites on lysosomes (Fig. 8C). With these parameters and conditions, the total fluorescence associated with individual lysosomes was converted to a corresponding number of V-ATPases, with the assumption that all three A subunits of every V-ATPase are accessible for SidK-AL568 binding. In HeLa cells, where individual lysosomes can be delineated more readily than in macrophages, the number of SidK-AL568 molecules associated per lysosome varied (Fig. 8D), with an average of 4.42 ± 0.03 (mean ± SEM) SidK molecules per lysosome, equivalent to 1.47 V-ATPase complexes per lysosome.

Previous research had shown that lysosomes are heterogeneous within a cell (Bright et al., 1997, 2016; Butor et al., 1995; Cheng et al., 2018). Additionally, lysosomal pH can vary with vesicle positioning, with peripheral lysosomes generally being more alkaline, and lysosomes near the center of the cell being more acidic (Johnson et al., 2016; Webb et al., 2021). A simple explanation for this finding could be that peripheral lysosomes have fewer V-ATPase complexes than central lysosomes, although other mechanisms could also cause this effect (see Introduction). To explore this phenomenon experimentally, lysosomes of HeLa cells were identified with LAMP1 as a marker and their V-ATPase density quantified with SidK-AL568. The resultant labeling with SidK-AL568 was more intense in juxtanuclear lysosomes than in more peripheral lysosomes (Fig. 8E). This differential distribution was quantified and is represented as a ratio of SidK-AL568:LAMP1 fluorescence (Fig. 8F). To more precisely assess whether the number of V-ATPases per lysosome varies as a function of their subcellular localization, the SidK-AL568 fluorescence of individual LAMP1-positive structures was analyzed relative to their distance from the edge of the cell. Cell outlines were drawn and degraded inward by 4 μm iteratively, to create concentric shells within each cell (Fig. 8G). Comparison of these subgrouped LAMP1-positive vesicles confirmed the existence of a gradient of SidK-AL568 that correlates with the distance of the lysosomes from the cell center. The most juxtanuclear lysosomes bind approximately twice as many SidK-AL568 molecules as peripheral ones (Fig. 8H). This change in V-ATPase density corresponded to 2.1 V-ATPase complexes per lysosome at > μm from the cell edge, compared to 1.1 lysosomal V-ATPases per lysosome for lysosomes 0 to 4 μm from the cell edge. Interestingly, ≈18% of the lysosomes closest to the nucleus showed more than the average number of V-ATPase complexes (2 to 6 per lysosome). We conclude that the pH heterogeneity of lysosomes within individual cells can be explained, at least in part, by differences in their V-ATPase density.

## DISCUSSION

Despite its clear importance in the maintenance of organellar pH and tissue homeostasis, the study of V-ATPases has been hampered by the lack of reagents to accurately localize and quantify these complexes. To address this problem, we developed a detection tool based on SidK, a *L. pneumophila* effector protein. SidK was an attractive detection tool for several reasons. First, it was reported to bind to the V-ATPase A subunit with high specificity (Abbas et al., 2020; Zhao et al., 2017; Xu et al., 2010) and affinity; SidK binds to the V-ATPase with a *K_d_* ≈ 3.5 nM (Sharma and Wilkens, 2017), which is comparable to the affinity of many immunoglobulins for their cognate antigens. Second, because the primary sequence of the V-ATPase A subunit is highly conserved across Eukarya, a SidK-based reagent should be applicable to a variety of model organisms. Indeed, we found SidK to interact equally with yeast and mammalian cells. Finally, unlike other V-ATPase subunits, the A subunit exists as a single isoform, so that the SidK reagent would be expected to bind V-ATPase wherever present in organelles, cells, and tissues.

We first expressed the V-ATPase-binding region of SidK (amino acids 1 to 278) as a fusion with GFP or mCherry. At modest expression levels, this construct enabled the detection of V-ATPase-rich membranes. However, at higher expression levels the unbound (cytosolic) probe obscured the intracellular organelles. This problem could be mitigated by leaching the cytosolic excess out the cells by permeabilizing the plasmalemma with PLY, revealing the specifically-bound constructs. However, when expressing SidK, a process that requires hours, we observed a change in lysosomal positioning that we attributed to its inhibitory effect, as it mimicked the effects of concanamycin A. Accordingly, we measured a ≈ 30% inhibition of the proton-pumping rate in cells expressing SidK-mCherry. While our findings were not unexpected considering the observations made earlier *in vitro* and in cells (Abbas et al., 2020; Zhao et al., 2017; Xu et al., 2010; Johnson et al., 2016), they nonetheless underscored a limitation inherent to the use of ectopically expressed SidK. Indeed, inhibition of V-ATPase and the resultant changes in luminal pH can affect not only lysosome localization, but also the ability of the V-ATPase to interact with other molecules (Maranda et al., 2001; Hurtado-Lorenzo et al., 2006; Hosokawa et al., 2013), altering intracellular signaling (Balgi et al., 2011; Hu et al., 2016). In this context, it is noteworthy that SidK-GFP/mCherry did not properly label acidic portions of the Golgi complex. This lack of labeling could have resulted from obstruction caused by other molecules that associate with V-ATPase in the Golgi, but not with those of the endocytic compartment.

To circumvent the problems associated with SidK expression, we utilized fluorescently-conjugated recombinant SidK as an overlay staining reagent, followed by washing to remove unbound probe. SidK-AL568 localized to punctate and cisternal structures in mammalian cells, with little nonspecific background. The high signal-to-noise ratio, along with specific organellar markers allowed us to clearly discern V-ATPase-positive organelles, primarily endo/lysosomes and the *trans*-Golgi. This labeling was in sharp contrast to the staining we obtained using a commercially available antibody to the A subunit and the conditions for its use described in the literature (Ramirez et al., 2019), which yielded results similar to those reported by others using other available antibodies (Yajima et al., 2007; McGuire et al., 2019; Michel et al., 2013). V-ATPase complexes have been shown to interact with several accessory proteins involved in endocytic sorting and signaling (Merkulova et al., 2015; Maxson and Grinstein, 2014; Marshansky et al., 2014). These interacting proteins include sorting nexins SNX27, SNX10 and SNX11 (Merkulova et al., 2015; Chen et al., 2012; Xu et al., 2020), and the Ragulator complex, which controls mTOR activation during nutrient sensing (Zoncu et al., 2011). Additionally, the V-ATPase is hypothesized to function as a sensor of luminal pH, that regulates endocytosis through a pH-dependent interaction with ARNO, the guanine nucleotide exchange factor for Arf6 (Hosokawa et al., 2013; Maranda et al., 2001; Hurtado-Lorenzo et al., 2006). Recently, some members of TLDc protein family have been found to interact with V-ATPase domains and modulate V-ATPase activity or stability (Merkulova et al., 2015; Castroflorio et al., 2021). These reports have relied on immunoprecipitation with available V-ATPase antibodies to study interactions. However, use of the more specific SidK-AL568 probe could provide more sensitive and/or specific measurements of interactions, as well as the ability to visualize novel interactions by high-resolution microscopy. Of interest, the 34.6 kDa SidK_1-278_ fragment we used is considerably smaller than conventional primary antibodies (IgG ≈150 kDa), which are often used in combination with an equally large secondary antibody. Our smaller probe is better suited to access V-ATPases, and could therefore be advantageous for immunogold-labeling for transmission electron microscopy, where the penetrance of bulky antibodies can limit detection. The smaller size of SidK-AL568 compared to SidK-GFP/mCherry may also explain why the former is better able to detect V-ATPase in the Golgi complex, although fixation and permeabilization may have also facilitated its access.

The successful labeling of the acidic intracellular compartments emboldened us to utilize the probe to monitor the dynamics of recruitment of the V-ATPase to a specialized vesicular compartment, the phagosome. The progressive development of phagosomal acidification is well established and was assumed, yet not proven, to be associated with increasing V-ATPase density. We found that while the nascent phagosome is virtually devoid of V-ATPases, the presence of the complex becomes evident in early (Rab5- and PtdIns(3)P-positive) phagosomes, and their number increases further as they attain the phagolysosomal stage. Therefore, the increasing acidity during maturation can be attributed, at least in part, to an increase in V-ATPase density. This increase in density correlates well with the recent finding that acidification of phagosomes to a pH < 6 during the transition between early and late phagosome results in the dissociation of Vps34 class III phosphatidylinositol-3-kinase from these organelles (Naufer et al., 2018). Therefore, V-ATPase recruitment to the phagosome would be expected to control the cessation of PtdIns(3)P synthesis on these compartments, which is necessary for transition to the late phagosome/phagolysosome stage (Vieira et al., 2001; Naufer et al., 2018).

We also observed that V-ATPase was not homogeneously localized throughout the membrane of the phagosome, and similar observations were made in enlarged and normal lysosomes. The existence of specialized subdomains was recently reported in phagosomes, where it was associated with the formation of contacts with the ER and the extension of tubular structures (Levin-Konigsberg et al., 2019). In the endocytic system, similar microdomains have been postulated to play roles in cargo sorting and receptor recycling through tubulation (Wijdeven et al., 2016; Rocha et al., 2009), and are likely also involved in phagosome resolution. Once internalized material has been degraded by acidic hydrolases, essential phagosomal and lysosomal membrane proteins must be parsed and redistributed within the cell for reutilization. The reformation of the lysosomal compartment by membrane-retrieval following content condensation was predicted long ago (Bright et al., 1997, 2016; Mullock et al., 1998), but the fate of specific components has not been well documented. In the case of the V-ATPase, retrieval from maturing phagosomes was demonstrated in *Dictyostelium discoideum* (Clarke et al., 2010), but to our knowledge has not been studied in mammalian cells. The observed V-ATPase-rich domains may be a precursor to the formation of recycling vesicles or tubules. In this regard, pH gradients have been recently observed in tubules formed by macrophages (Suresh et al., 2020; Naufer et al., 2018). Taken together, these findings suggest that proton-pumping microdomains may form on acidic organelles during the tubulation and fission that accompany resolution.

The high signal-to-noise ratio provided by the SidK-AL568 probe and its defined labeling stoichiometry enabled us to quantify the number of V-ATPase complexes per organelle. Quantification relied on the use of saturating concentrations of the probe, a requirement that was fulfilled by analyzing the concentration dependence of SidK-AL568 binding. We calculated the number of V-ATPases per lysosome to range between 1 and 6, averaging ≈1.5. This number is similar to estimates for synaptic vesicles, with one V-ATPase per vesicle (Takamori et al., 2006). The paucity of pumps per organelle likely accounts for the difficulty encountered in their detection and highlights the need for highly-specific probes. It is important to emphasize that our estimates for the number of V-ATPases rely on the assumption that all three A subunits of every pump are accessible to the probe. Because an increasing number of proteins are appreciated to bind to the V-ATPase, steric hindrance may curtail the number of available subunits, which would result in an underestimate of the number of pumps. Through these experiments, we noted that the distribution of V-ATPases in association with LAMP1-positive structures was not homogeneous throughout the cell: more SidK-AL568 bound to juxtanuclear lysosomes, when compared to peripheral lysosomes. Shell analysis showed that the density of SidK-AL568 increased with distance from the cell edge, with the most central lysosomes containing up to 6 V-ATPases. These data are in line with recent reports showing that juxtanuclear lysosomes are, on average, more acidic than the more peripheral ones (Johnson et al., 2016; Webb et al., 2021).

While bearing in mind the caveats raised above regarding the assumption of unimpeded access to all the A subunits, our estimates can be used to calculate the rate of pumping of individual lysosomal V-ATPases *in situ*. Experiments like those in Fig. 2H indicated that addition of concanamycin alkalinized the lysosomes at an initial rate of 0.198 ± 0.1 pH.min^-1^. Because at steady state the activity of the pumps must have matched the proton leak unmasked by concanamycin, the measured rate can be used to estimate pump activity. Considering the buffering power of the lysosomes, which we estimated to be 23.5 mM.pH^-1^ –measured by pulsing with ammonium– we calculate a proton flux per lysosome volume of 9.50 × 10^-2^ mmol.min^-1^.L^-1^. Assuming an approximate lysosomal volume of 2 × 10^-17^.L^-1^ (de Araujo et al., 2020; Yordanov et al., 2019), this measurement is equivalent to a flux of 953 H^+^.sec^-1^ per lysosome. As ten protons are pumped by the V-ATPase for every three ATPs hydrolyzed (Zhao et al., 2015), the flux corresponds to approximately 191 molecules of ATP hydrolyzed per lysosome per sec. Considering that there are 1 to 2 pumps on average per lysosome, the calculated flux is within the same order as the V-ATPase activity measured previously in cell-free systems in mammals (29.2 ATP per V-ATPase.sec^-1^; Abbas et al., 2020) and yeast (from 3.7-300 ATP per V-ATPase.sec^-1^; Vasanthakumar et al., 2019; Uchida et al., 1985; Sharma and Wilkens, 2017; Kawasaki-Nishi, 2001). The rate estimated here in intact cells may differ from some of the *in vitro* measurements due to the effect of regulatory factors and parameters, such as availability of counterions, phosphorylation, membrane lipid composition, etc. that are not encountered in cell-free enzyme assays.

In summary, we have introduced a new, powerful probe to visualize and quantify V-ATPases in eukaryotic cells. This probe enabled us to assess the subcellular distribution of V-ATPases and, in combination with ratiometric pH determinations, estimate the rate of flux for individual lysosome-associated V-ATPase complexes in intact cells. These results reveal heterogeneity in the lysosomal compartment that is a result of vesicle location dependent differences in V-ATPase density. We also expect this probe to be a useful label for super-resolution imaging and electron microscopy, and anticipate that SidK-AL568 and similar SidK-derived probes will contribute not only to studies of V-ATPase complexes in cells, but to our understanding of endocytic and secretory processes more generally.

## MATERIALS AND METHODS

### Cell culture

HeLa and RAW264.7 cells were obtained from and authenticated by the American Type Culture Collection (ATCC). Both cell lines tested negative for mycoplasma contamination by DAPI staining. HeLa cells were grown in DMEM containing L-glutamine and 10% heat-inactivated fetal calf serum (FCS; MultiCell, Wisent) at 37°C under 5% CO_2_. RAW264.7 cells were grown in RPMI-1640 medium containing L-glutamine and 10% heat-inactivated FCS, at 37°C under 5% CO_2_.

To obtain non-polarized human monocyte-derived macrophages (M0 hMDMs), peripheral blood mononuclear cells (PBMCs) were isolated from the blood of healthy donors by density-gradient separation with Lympholyte-H (Cedarlane). Human monocytes were then separated by adherence and incubated in RPMI-1640 containing L-glutamine, 10% heat-inactivated FCS, 100 U.mL^-1^ penicillin, 100 μg.mL^-1^ streptomycin, 250 ng.mL^-1^ amphotericin B and 25 ng.mL^-1^ hM-CSF (PeproTech) for 5 to 7 days, at 37°C under 5% CO_2_, before experimentation.

### Reagents

Mammalian expression vectors were obtained from the following sources: pmCherry-N1 (Clontech) and pEGFP-N1 (Clontech), sec61b-GFP (Addgene; plasmid no. 121159), PM-GFP (Teruel et al., 1999), Rab5A-GFP (Roberts et al., 2000), PX-GFP (Kanai et al., 2001), Rab7-GFP (Bucci et al., 2000), LAMP1-GFP (Martinez et al., 2000), ORP1L-GFP (Rocha et al., 2009), Arl8b-GFP (Johnson et al., 2016). V_O_a1-, V_O_a2-, and V_O_a3-GFP were the kind gift of Dr. Shuzo Sugita (UHN, Toronto).

Primary antibodies were purchased from the following vendors: anti-human LAMP1 (Developmental Studies Hybridoma Bank; catalogue no. H4A3-s), anti-VapB (Sigma-Aldrich; catalogue no. HPA013144), anti-TGN46 (Abcam; catalogue no. ab50595), anti-GM130 (BD Transduction Laboratories; catalogue no. 610822), anti-V1A (Abnova; catalogue no. H00000523-M02). Secondary antibodies conjugated with Alexa Fluor-405, -488, -555 or -647 were purchased from Jackson ImmunoResearch Labs.

Fluorescently-conjugated 10 kDa dextrans, Alexa Fluor-405 and Alexa Fluor-568 NHS esters, and fluorescently conjugated phalloidin and streptavidin were purchased from Invitrogen. Concanamycin A, nigericin, DAPI, propidium iodide and cresyl violet were from Sigma-Aldrich. Sheep red blood cells (SRBC; 10% suspension) were from MP Biomedicals. Anti-sheep red blood cell antibodies were from Cedarlane Laboratories. Paraformaldehyde (PFA; 16% wt/vol) was from Electron Microscopy Sciences.

### SidK-GFP/mCherry plasmid construction

SidK_1-278_ was amplified by PCR using plasmid pSAB35 (Abbas et al., 2020) as a template, with the following forward and reverse primers, respectively: 5’-GAGGAGGAATTCATGTCTTTTATCAAGGTAGGTATAAAAATG-3’ and 5’-GAGGAGGGATCCCCTTTGCTTAAAGCATTTAATTTTTCG-3’. The PCR product was digested with EcoRI and BamHI (New England Biolabs), and ligated into pmCherry-N1 and pEGFP-N1 plasmids that had been digested with the same restriction enzymes.

For illustrative purposes, a model of yeast V-ATPase with SidK-GFP bound (Fig. 1B) was generated from PDB accession no. 5VOX, 1GFL and 6O7T, using *UCSF ChimeraX* (Goddard et al., 2018).

### Purification of SidK_1-278_ protein

To purify SidK_1-278_-3×FLAG, *Escherichia coli* strain BL21 was transformed with pSAB35 (Abbas et al., 2020) and grown at 30°C with shaking in 1 L LB medium (BioShop) supplemented with 50 mg.L^-1^ kanamycin. At an OD_600_ of 0.6-0.8, protein expression was induced with 1 mM IPTG and cells grown overnight at 16°C. All subsequent steps were performed at 4°C. Cells were harvested by centrifugation at 5,250 ×*g*, resuspended in 25 mL HisTrap Buffer (50 mM Tris-HCl pH 7.4, 25 mM imidazole, and 300 mM NaCl) and lysed by sonication. The cell lysate was centrifuged at 38,000 ×*g* and the supernatant was loaded onto a 5 mL HisTrap Ni-NTA column (GE Healthcare). The column was washed with HisTrap Buffer and protein eluted with a linear gradient of imidazole from 25 to 300 mM in HisTrap Buffer over 10 column volumes.

Fractions containing 6×his-SidK_1-278_-3×FLAG were pooled, mixed with TEV protease, and dialyzed against 2 L Dialysis Buffer (50 mM Tris-HCl pH 7.4 and 300 mM NaCl) with 1 mM dithiothreitol (DTT) overnight. Cleaved protein was dialyzed against 2×1 L Dialysis Buffer to remove imidazole and DTT and passed through a 5 mL HisTrap column. The column was washed with HisTrap Buffer and the flowthrough and wash were collected, pooled, and concentrated in a centrifugal concentrating device (EMD Millipore). To remove aggregated protein, SidK_1-278_-3×FLAG was further purified with a Superdex 200 10/300 Increase gel filtration column (GE Healthcare) equilibrated with 50 mM Tris-HCl pH 7.4 and 150 mM NaCl. Fractions containing protein were pooled, concentrated, flash-frozen in liquid N_2_, and stored at -80°C.

### Purification of *Saccharomyces cerevisiae* V-ATPase using SidK

*Saccharomyces cerevisiae* strain BJ2168 was grown in 11 L yeast extract peptone dextrose medium (YPD; BioShop) in a Microferm fermenter (New Brunswick Scientific) at 30°C, with aeration of 34 cubic feet per hour, and stirring at 300 rpm. Yeast were harvested after 18 h (OD_660_ = 4.5) by centrifugation at 4,000 x*g* for 15 min at 4°C. All subsequent steps were performed at 4°C. Cell walls were broken by bead beating in lysis buffer (phosphate-buffered saline, pH 7.4, 8% (w/v) sucrose, 2% (w/v) sorbitol, 2% (w/v) glucose, 5 mM □-aminocaproic acid, 5 mM *p*-aminobenzoic acid, 5 mM EDTA, and 0.001% (w/v) PMSF). Cellular debris was removed by centrifugation at 3,000 x*g* for 10 min and cell membranes were collected by ultracentrifugation at 152,957 x*g* for 40 min. The membrane pellet was resuspended in 36 mL lysis buffer, divided into 4 aliquots, flash-frozen in liquid N_2_, and stored at -80°C. Two membrane pellets (corresponding to half a fermenter growth) were thawed and solubilized with addition of n-dodecyl-β-D-maltopyranoside (DDM, Anatrace) to 1% (w/v) final concentration, and DDM-solubilized *S. cerevisiae* V-ATPase isolated with M2 Affinity agarose gel (Sigma-Aldrich) pre-loaded with SidK_1-278_-3×FLAG as described previously (Abbas et al., 2020). Protein purity was confirmed by SDS-PAGE using 4–20% Mini-PROTEAN TGX protein Gels (BioRad).

### Transient DNA Transfection

HeLa cells were plated at on 18 mm glass coverslips at a concentration of ≈5×10^4^ cells.mL^-1^, 16-24 hr prior to transfection. FuGENE 6 (Promega) transfection reagent was used according to the manufacturer’s instructions to transfect HeLa cells at a 3:1 ratio (using 1.5 μL FuGENE 6 and 0.5 μg DNA per well). RAW264.7 cells were plated on 18 mm glass coverslips at ≈2×10^5^ cells.mL^-1^, 16-24 hr prior to transfection. FuGENE HD (Promega) transfection reagent was used to transfect RAW264.7 cells at a 3.5:1 ratio (using 1.75 μL FuGENE HD and 0.5 μg DNA per well). In all cases, monolayers were used for experiments 16 hr after transfection.

### Ratiometric fluorescence microscopy for the measurement of lysosomal pH, V-ATPase activity and buffering power

HeLa cells were plated on 18 mm glass coverslips and incubated overnight with 250 μ fluorescein isothiocyanate (FITC)-conjugated 10 kDa-dextran, which was then chased for 1 h in complete medium prior to imaging to visualize lysosomes. Coverslips were then mounted in a Chamlide magnetic chamber and incubated in HBSS medium for fluorescence-based pH determinations.

Steady-state lysosomal pH was determined by exciting FITC-dextran labelled lysosomes sequentially at 481±15 nm and 436± 20 nm, collecting emitted light at 520±35 nm. The fluorescence intensity of FITC when excited at ≈490 nm is highly pH dependent and was used to determine the pH of lysosomes. The fluorescence when excited at ≈440 nm is much less pH dependent and was used to correct for potential photobleaching or focal changes during image acquisition, the 490 nm/440 nm fluorescence ratio of FITC is utilized to determine pH values. Multiple fields of cells were imaged, and the data processed with Volocity.

The 490 nm/440 nm fluorescence ratios were converted to pH by sequentially incubating the cells for 5 min in isotonic K^+^ solutions (143 mM KCl, 5 mM glucose, 1 mM MgCl_2_, 1 mM CaCl_2_) of different pH (pH 4.5, buffered with 20 mM acetic acid, pH 5.5, 6.5, 7.5 solutions buffered with 20 mM MES), containing 10 μM nigericin and 5 μM monensin. Background-subtracted 490/440 nm fluorescence ratios were then plotted against pH, and the data fitted by least squares were used to interpolate the lysosomal pH.

For V-ATPase activity determinations, HeLa cells were acutely treated with 500 nM concanamycin A and images were acquired at 30 s intervals for 10 min. The time dependence of the resulting pH changes was then plotted in GraphPad Prism 6, and the initial rates of alkalinization (per cell) were determined within 2 min of addition of concanamycin A.

For the determination of buffering power, HeLa cells containing FITC-dextran loaded lysosomes were challenged with 0.5 mM NH_4_Cl and the resulting change in fluorescence intensity (490 nm/440 nm) was measured immediately. The corresponding pH change, calculated as above, was used to calculate the concentration of NH_4_ formed inside the lysosome, using the Henderson-Hasselbalch equation, and the buffering capacity estimated as Δ[NH_4_^+^]_lys_/ΔpH_lys_.

### Lysosome labeling, cresyl violet staining and dissipation of organellar pH

For identification of lysosomes, HeLa cells were incubated overnight with 100 μg.mL^-1^ fluorescently-conjugated 10 kDa-dextran at the time of transient transfection, 16 to 24 hr prior to experiments. The next day, monolayers were washed 3× with PBS and placed in HBSS medium at 37°C for 1 hr to chase the dextran to lysosomes. Monolayers were then imaged live by confocal microscopy.

For cresyl violet staining, HeLa cells, seeded on 18 mm glass coverslips at a concentration of 1×10^5^ cells.mL^-1^, were labeled with fluorescent dextran as described above. Monolayers were then incubated at 37°C with 1□μM cresyl violet in Hanks’ Balanced Salt Solution (HBSS; MultiCell, Wisent) for 5 □min, washed 3 times and imaged live by confocal microscopy. In some cases, organellar pH was dissipated prior to cresyl violet staining. To this end the cells were treated for 30□min with 250 nM concanamycin A and 10□mM NH_4_Cl in HBSS. After this treatment, monolayers were stained with 1 μM cresyl violet and imaged live, as above.

### Fluorescent labeling of SidK_1-278_

Prior to labeling, purified SidK_1-278_-3×FLAG (see above; referred to in the text as SidK) was buffer exchanged into PBS using a centrifugal concentrator (EMD Millipore). Following this, SidK was directly labeled by conjugation with Alexa Fluor-568 NHS-ester (Invitrogen). A 10:1 dye:SidK molar ratio was prepared in 0.05 M borate buffer, vortexed, and incubated shaking at 500 rpm for 1 hr at room temperature. Labeled SidK (referred to as SidK-AL568) was then dialyzed in 4× 1 L PBS, to remove unincorporated dye. One volume of 100% glycerol was added to the SidK-AL568 for stability, and stored at 4°C.

The fraction of SidK labeled with Alexa Fluor-568 was determined using absorbance spectroscopy on a Nanodrop 2000 instrument (Thermo Scientific). SidK and SidK-AL568 was diluted to 0.1 mg.mL^-1^ in PBS. The *A*280 of SidK and its molecular weight (34,646 g.mol^-1^) was used to calculate its extinction coefficient (ε *SidK* = 41,690 cm^-1^.M^-1^). Then, the *A*280 and *A*577 (λ_max_ of Alexa Fluor 568) of SidK-AL568 were measured and the degree of labeling (moles of dye per mole of protein) for SidK-AL568 calculated, using the following two equations, along with the *ε dye* value for Alexa Fluor 568 (91,300 cm^-1^.M^-1^) and the correction factor for the *A*280 contribution of Alexa Fluor-568 (*CF*280 = 0.46):

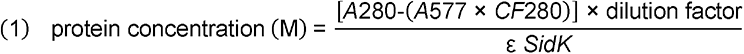

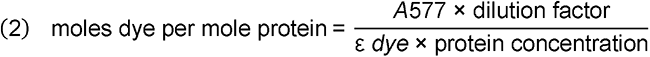

The number of dye particles per SidK-AL568 molecule was determined by single-molecule photobleaching imaging (Liesche et al., 2015). Briefly, a 10^6^ dilution of SidK-AL568 was prepared and added to clean coverslips, creating a monodispersed sample. Monodispersed SidK-AL568 was then imaged continuously by confocal microscopy at ≈ 5 frames.sec, until spot disappearance occurred through photobleaching. Time series images were analyzed using Volocity v6.3 software (Quorum Technologies), and SidK-AL568 molecule fluorescence over time plotted to generate traces where the number of steps required for SidK-AL568 spot disappearance were counted. Each step corresponded to a single Alexa 568 dye molecule conjugated to SidK. The average fluorescence value for *bona fide* single bleach step SidK-AL568 was used to calculate the percent of SidK-AL568 in the monodispersed population with 1, 2, 3, or 4 or 5+ dye molecules per SidK.

### Culturing of yeast and immunofluorescence of yeast spheroplasts

Wild type (*MAT*α *ura3–52 leu2–3,112 his4–519 ade6)* and *vma1*Δ *(MAT* α *ura3–52 leu2–3,112 his4–519 ade6 vma1*Δ *::LEU2*) *S. cerevisiae* strains were kindly gifted by Dr. Morris Manolson (University of Toronto). Yeast cultures were routinely grown at 30^°^C, with shaking, in YPD medium (BD Biosciences).

For spheroplast preparation, overnight cultures were diluted 1:100 in YPD broth and grown at 30^°^C for 2 hr. One mL of yeast culture was then fixed with 4% PFA for 30 min, while shaking at room temperature. After washing cells 2× with 0.1 M KHPO_4_ pH 6.5, yeast were washed 1× in K-Sorb (1.2 M sorbitol in 0.1 M KPHO_4_) and resuspended in 0.5 mL K-Sorb containing 5 μL β-mercaptoethanol and 10 mg.mL^-1^ zymolyase 20T (MP Biomedicals). Cells were shaken at room temperature for 30 min, and gently washed 2× with K-Sorb. After final resuspension in 0.5 mL K-Sorb, 20 μL of spheroplast suspension was spread on a concanavalin A-coated slide and allowed to adhere for 15 min at room temperature. Following this, spheroplasts were permeabilized/blocked for 30 min in PBS 0.1% Tween-20 containing 5% BSA and 5% skim milk. Samples were then incubated in the same buffer containing 1:200 SidK-AL568 and 1 μg.mL^-1^ DAPI for 30 min at room temperature. After staining, slides were washed 3× with PBS and visualized by confocal microscopy.

### Immunofluorescence of mammalian cells

After fixing in 3% PFA for 10 min at room temperature, cells were washed in PBS and permeabilized/blocked in PBS containing 5% BSA and 0.1% Triton X-100 for 30 min at room temperature. Samples were then incubated with primary staining reagents for 30 min at room temperature. Primary antibody dilutions were: LAMP1 (1:50), VapB (1:100), TGN46 (1:100), GM130 (1:100), V1A (1:100). For mitochondrial staining, fluorescently-conjugated streptavidin was used at 1:100 dilution. SidK-AL568 was used at 1:100, 1:50 or 1:25 dilution, as indicated in the text. Where indicated, a 5-fold excess of unlabeled SidK_1-278_ was used in the blocking/permeabilization step, to block SidK-AL568 binding sites prior to the addition of SidK-AL568. After primary staining, monolayers were washed 3× with PBS, and samples incubated 30 min at room temperature with Alexa Fluor-conjugated secondary antibodies at a 1:1000 dilution. Where indicated, 1:1000 fluorescent phalloidin or 1μg.mL^-1^ DAPI was added together with the secondary antibodies. Samples were washed 3× with PBS and viewed by confocal microscopy in PBS.

### Phagosome and sucrosome analyses

For phagocytosis assays, RAW264.7 cells were plated at on 18 mm glass coverslips at a concentration of 2×10^5^ cells.mL^-1^ and grown for 16-24 hr. Cells were transfected as described above with the constructs indicated in the text. The day of experiments, SRBCs were prepared for use in phagocytosis. Briefly, 100 μL of SRBC suspension was washed with 3× with PBS and labeled with Alexa Fluor-405 NHS ester for 20 min, shaking at room temperature. SRBC were then opsonized with 2 μL of rabbit anti-SRBC IgG at 37°C for 1 h. Prepared SRBCs were washed 3× with PBS and resuspended to a final volume of 1 mL in PBS. After 1:10 dilution in PBS, 25μL of this suspension was added to the RAW264.7 cells. Alternatively, FITC-labelled zymosan was diluted to 10 mg.mL^-1^ in PBS and opsonized by incubation with human IgG (final IgG concentration, 5 mg.mL^-1^) for 30 min at room temperature. Prepared zymosan was then washed 3× with PBS and resuspended to a final volume of 20 µL in PBS. 1 μL of this suspension was added to RAW264.7 seeded onto coverslips. In all cases, phagocytosis was synchronized by sedimenting particles onto the cells using centrifugation at 300 ×*g* for 1 min. After phagocytosis, monolayers were fixed in 3% PFA for 10 min at room temperature and stored in PBS until used. FITC-zymosan phagocytosis, which was used to determine the rate of acidification of the nascent phagosome, was imaged immediately after sedimentation of the particles.

To generate sucrosomes, transfected RAW264.7 cells were incubated for 16 to 24 hr in RPMI-1640 medium containing L-glutamine, 10% heat-inactivated FCS and 30 mM sucrose, at 37°C under 5% CO_2_. The next day, cells were washed 3× with PBS, placed in RPMI-1640 medium containing L-glutamine and 10% heat-inactivated FCS, and used for experiments as described in the text. After experiments, monolayers were fixed in 3% PFA for 10 min at room temperature and stored in PBS.

### Quantitation of the number of SidK-AL568 per lysosome

The average fluorescence value of monodispersed SidK-AL568 molecules was used to estimate the number of SidK-AL568 monomers per lysosome in HeLa cells that had been co-stained with SidK-AL568 and LAMP1. 1:25, 1:50 and 1:100 SidK-AL568 dilutions were assessed to determine the conditions required for saturation of SidK-AL568 staining, needed to estimate the maximum number of SidK-AL568 molecules bound per LAMP1^+^ lysosome.

### Microscopy

Confocal images were acquired using a spinning disk system (Quorum Technologies Inc.). The instrument consists of a microscope (Axiovert 200M; Zeiss), scanning unit (CSU10; Yokogawa Electric Corporation), electron-multiplied charge-coupled device camera (C9100-13; Hamamatsu Photonics), five-line (405-, 443-, 491-, 561-, and 655-nm) laser module (Spectral Applied Research), and filter wheel (MAC5000; Ludl) and is operated by Volocity v6.3. Images were acquired using a 63×/1.4 NA oil objective (Zeiss), with an additional 1.5× magnifying lens and the appropriate emission filter. For live experiments, cells were maintained at 37°C using an environmental chamber (Live Cell Instruments).

Ratiometric fluorescence pH measurements were acquired on an epifluorescence microscope (Axiovert 200M; Zeiss) running on Volocity v6.3, equipped with a camera (Flash 4.0v2; Hamamatsu), excitation lamp (X-Cite 120; EXFO Life Sciences Group), a 63x/1.4 NA oil objective (Zeiss), the appropriate dichroic mirror (CFP/YFP; Chroma Technology), and filter wheels containing the necessary filters for FITC ratiometric fluorescence determinations (excitation at 481±15 nm or 436±20 nm, emission 520±35 nm). All experiments were performed maintaining the temperature at 37°C with an environmental controller (Medical Systems Corporation).

### Image analysis

Image processing and analyses were performed using Volocity v6.3. Image deconvolution was done on acquired Z-stacks within the Volocity Restoration module, using the iterative restoration function. Calculated fluorochrome point spread functions were used to deconvolve individual channels for 5-8 iterations, until a confidence limit of >90% was achieved. For colocalization analyses, Volocity Colocalization module was used to calculate the positive product of the differences of the mean channels (Li, 2004), which was then overlaid on merged images for visualization. Alternatively, for some colocalization analyses, Manders’ overlap coefficients were calculated in Volocity, which describe the percent of various organelle markers that colocalize with SidK-AL568 (referred to as M).

The Volocity Ratio function was used for SidK-AL568 ratio calculations. This divides background-subtracted intensities of SidK-AL568 and lysosomal, sucrosomal or phagosomal markers (as indicated in the text), to calculate the ratio of SidK-AL568 fluorescence to that of the chosen organellar marker. The Ratio function generates a rainbow LUT ratio channel that is applied as an overlay on merged images, with a scale representing ratio values from 0 to 2.

### General methodology and statistics

Data calculations and normalizations were done using Microsoft Excel 2011 (Microsoft Corporation) or GraphPad Prism v9 software (GraphPad Software, Inc.). Because experiments were, for the most part, *in vitro* imaging determinations of individual cells, samples were assigned to groups according to specific experimental treatments (control vs. experimental group). The number of individual experiments and the number of determinations per experiment were selected to attain an estimate of the variance compatible with the statistical tests used, primarily student’s *t* test. Each type of experiment was performed a minimum of three separate times (biological replicates) and a minimum of ten individual event determinations (equivalent, but not identical to technical replicates). Data was tested for normality, and appropriate testing applied. No data was excluded as outliers. All statistics were calculated using GraphPad Prism v9.

## Data availability

Experimental datasets that support the findings of this study are available from the corresponding authors upon reasonable request.

## Online supplemental material

**Figure S1** shows the (**A**) SidK-AL568 or (**B**) α-V1A staining of HeLa cells before (left) and after (right) the thresholding used in the colocalization analyses (Costes et al., 2004) of Figure 4C and 4H, respectively.

---

**Figure S1.**
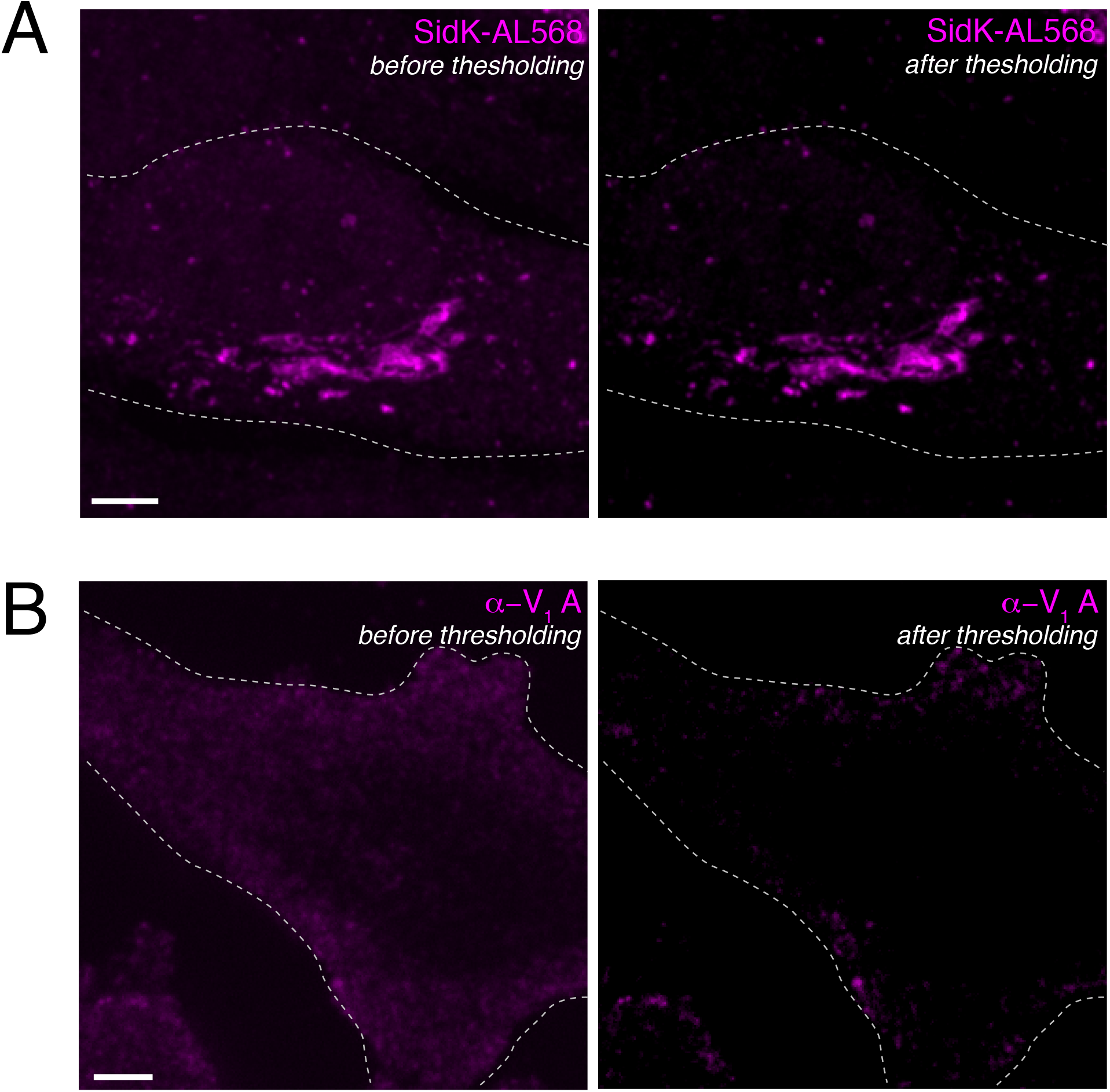
Demonstration of the Costes thresholding method used prior to colocalization analyses. HeLa cells were stained with (**A**) SidK-AL568 (magenta) or (**B**) α-V_1_A (magenta), as described in Materials and Methods. For both **A** and **B**, left panel shows staining before Costes thresholding, while the right panel shows the same channel after thresholding. Outlines of cells are indicated by dotted lines. Scale bars: 5 μm. Images in **A** and **B** correspond to Figure 4C and 4H, respectively.

